# Generative Design of Cell Type-Specific RNA Splicing Elements for Programmable Gene Regulation

**DOI:** 10.1101/2025.11.05.686847

**Authors:** Xi Dawn Chen, Maile Jim, Mounica Vallurupalli, Kai Cao, Andrea Navarro Torres, Jing Wesley Leong, Yifan Zhang, David Wollensak, Qiyu Gong, Jing Sun, Mehdi Borji, Gail Schor, Sofia Mrowka, Margaret Hu, Anisha Laumas, Jennifer A. Roth, Todd Golub, Fei Chen

## Abstract

Programmable control of gene expression in specific cell types is essential for both basic discovery and therapeutic intervention, yet current strategies lack scalability across diverse cellular contexts. Here, we introduce SPICE (Splicing Proportions In Cell types), an integrated experimental and computational framework that harnesses alternative RNA splicing as a programmable modality for cell type-specific gene regulation. To power SPICE, we constructed a massively parallel reporter assay (MPRA) comprising 46,372 human-derived sequences and profiled exon skipping across 43 cell lines spanning 10 lineages, uncovering widespread cell type-specific exon skipping. Using this data, we trained deep learning models that both predict splicing in unseen contexts and generate synthetic sequences with programmed, cell type-specific splicing patterns. Leveraging these models, we further engineered sequences that selectively splice in cells harboring oncogenic splicing factor mutations, demonstrating translational potential. SPICE provides a generalizable strategy for dissecting splicing regulation and engineering alternative splicing as a gene expression regulatory layer for research and therapeutic applications.

**One Sentence Summary:** We introduce SPICE, an integrated framework that couples large-scale splicing assays with generative design to uncover regulatory principles and design programmable, cell-specific gene expression for research and therapeutic applications.

## Introduction

Precise control of gene regulation remains a central challenge in both basic biology and therapeutic engineering. Achieving cell type-specific expression is critical for dissecting the functions of distinct cell populations and for enabling selective reporters, CRISPR-based therapeutics, and gene replacement therapies. Existing approaches, including nanoparticle targeting, viral vector engineering, *cis*-regulatory element (CRE)-based, and transcript sensing-based methods^1–5^, have yielded important successes but remain limited to particular tissues or cell states. Expanding this toolbox by harnessing additional levels of gene regulatory control would enhance the precision, efficacy, and safety of genetic therapies aimed at treating a broad spectrum of human diseases.

Alternative RNA splicing is a powerful but underexplored strategy for achieving programmable control of cell type-specificity. This process generates transcript and protein diversity through the selective inclusion or exclusion of specific exons and introns within pre-mRNA transcripts.

Splicing decisions are tightly regulated by a combination of splicing regulatory elements, most notably the core elements that include 5′ splice donor, 3′ splice acceptor, branchpoint, and polypyrimidine tract; as well as auxiliary elements like intronic and exonic splicing enhancers and silencers^6,7^. Over 95% of human multi-exon genes undergo alternative splicing to produce functionally distinct RNA isoforms, and aberrant splicing has been implicated in more than 15% of human genetic diseases and cancers^8,9^.

Alternative splicing has recently emerged as a promising strategy for driving cell type-or cell state-specific gene expression in therapeutic and biotechnological applications. While existing tools have demonstrated that splicing regulatory elements can be engineered to restrict gene expression to specific cellular contexts^10,11^, these efforts remain limited to narrow use cases and lack a generalizable framework for discovering and designing splicing regulatory elements across diverse cellular environments.

In parallel, advances in machine learning have shown that coupling large-scale datasets with predictive and generative models enables the design of novel cell type-specific CREs^4^. Yet analogous approaches for alternative splicing have lagged. While machine learning models have been trained to predict splice site usage, most remain cell type-agnostic and have not yet been extended to generative sequence design for programmable splicing control across cellular contexts^12–19^. This is largely due to the lack of proper training data: RNA-seq approaches enable transcriptome-wide splicing analysis across cell types but suffer from limited splice junction coverage and bias from gene expression differences^20–22^. Massively parallel reporter assays (MPRA) allow large-scale functional screening but have been mostly confined to single cell types^23–29^. Hence, we need both large-scale data generation for splicing across cell types and also novel machine learning models applied to such data.

Here, we introduce SPICE (Splicing Proportions In Cell types), an integrated experimental and computational framework for the systematic analysis and design of cell type-specific splicing sequences (Fig. 1a-c). SPICE combines: (1) large-scale MPRA profiling of 46,372 exon skipping events across 43 cell lines spanning 10 lineages from the Cancer Cell Line Encyclopedia (CCLE) (Fig. 1b), (2) Soma, a deep learning model that predicts splicing outcomes in unseen sequences and cell types, and (3) Melange, a generative model that designs synthetic sequences with specified cell type-specific splicing patterns (Fig. 1c). This dataset revealed hundreds of exons with robust cell type-specific splicing activity that can be harnessed to drive cell type-specific protein expression. Together, Soma and Melange enable both interpretation and design, decoding the sequence grammar of splicing specificity while generating novel sequences with tailored activity patterns. As a proof of concept, we applied SPICE to engineer sequences specific to splicing factor mutations that are common to many cancers. SPICE provides a unified framework for decoding the sequence grammar of cell type-specific splicing and for engineering synthetic sequences to precisely control gene expression.

**Figure 1:**
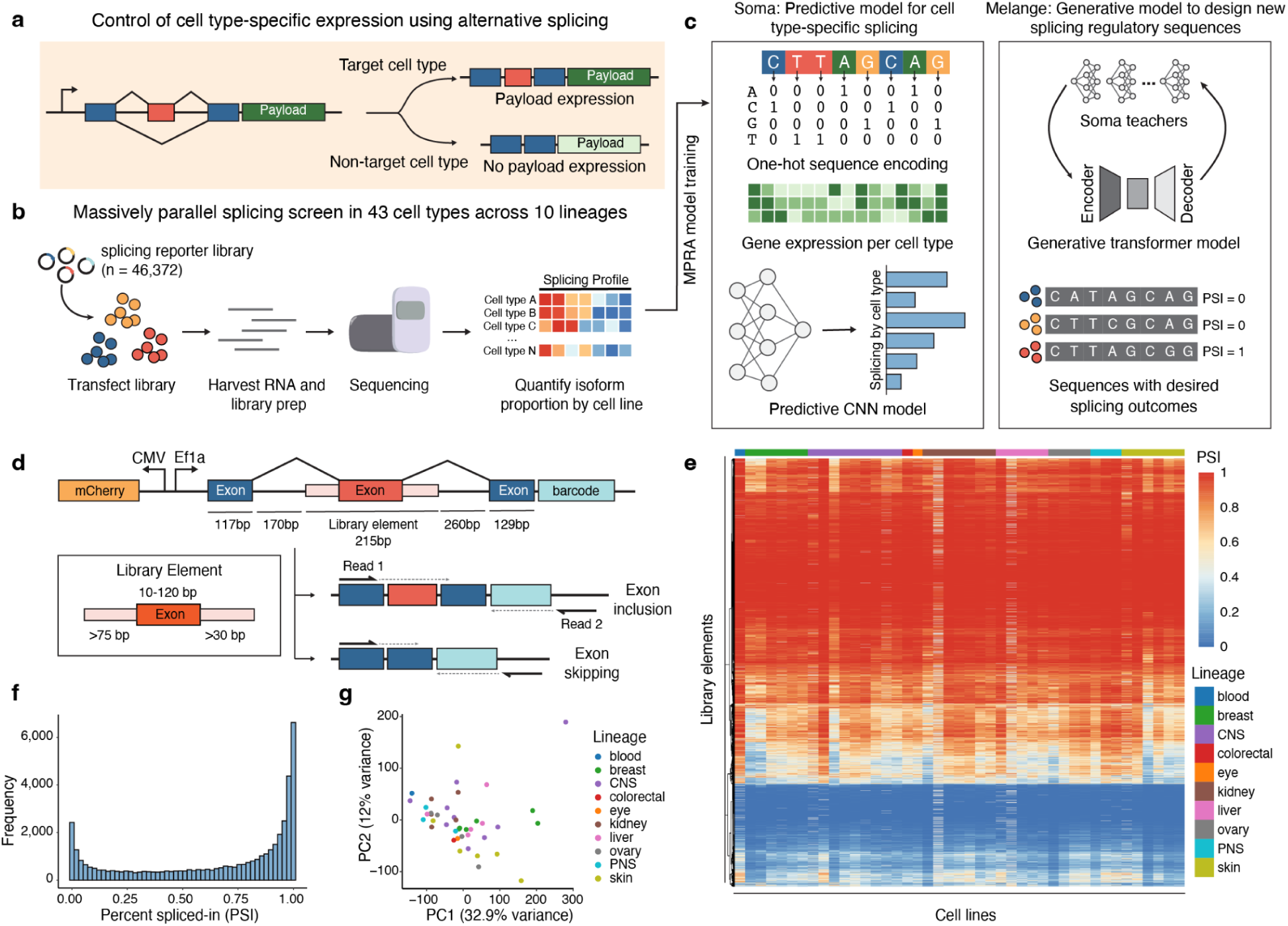
SPICE: an integrated experimental and computational framework for the systematic analysis and design of cell type-specific alternative splicing sequences. **a**, Conceptual overview using the SPICE framework to achieve cell type-specific gene expression. A cell type-specific alternative exon is placed upstream of a genetic payload to create two possible translational reading frames. The payload is expressed only in the target cell type when exon inclusion (or exclusion) produces a transcript that enables in-frame translation. **b**, Massively parallel splicing reporter screen across 43 cell types from 10 lineages. A pooled minigene library (n = 46,372 exons) is introduced into each cell line, RNA is harvested, and isoform proportions are quantified by sequencing. **c**, Overview of computational models. Soma is a predictive convolutional neural network trained on MPRA splicing data and cell type gene expression, while Melange is a generative variational autoencoder guided by Soma teachers to design new splicing elements with specified outcomes. **d**, Reporter design. Library elements, comprising synthesizable genomic exons with native flanking intron sequences, are inserted between constant upstream and downstream exons. Following transcription and splicing, the reporter yields either an exon-included or exon-skipped isoform. The reporter amplicon is sequenced, with read 1 spanning exon-exon junctions and read 2 capturing the barcode information. **e**, Heatmap of percent spliced-in (PSI) values for all library elements across 43 cell lines. Cell lines (x-axis) are clustered by lineage, and library elements (y-axis) are clustered by splicing outcome. Heatmap color scale indicates PSI, where red represents PSI = 1, and blue represents PSI = 0. **f**, Distribution of mean PSI values across all library elements, showing a bimodal distribution with most exons either highly included or highly skipped. **g**, Principal component analysis (PCA) of splicing profiles across cell lines. Points are colored by annotated CCLE lineage.

## Results

### Design of the splicing MPRA reporter library

To generate a large-scale dataset of alternative splicing activity across diverse cell types, we required a controlled, scalable reporter that decouples splicing measurements from endogenous gene expression. We developed a minigene system prioritizing exon skipping, the most frequent alternative splicing event^20^. Each reporter contained a variable library exon flanked by constant upstream and downstream exons, along with fragments of the native intronic contexts (Fig. 1d). Each minigene transcript also included a unique 14-nucleotide (nt) barcode in its 3′ untranslated region (UTR) for isoform identification, allowing recovery of the library sequence even when the exon was skipped. To capture splicing outcomes at high resolution, we used paired-end sequencing, with Read 1 spanning the splice junctions and Read 2 reading through the unique molecular identifier-(UMI) tagged 3′ UTR barcode (Supplementary Table 1). In our pilot experiments, we tested three reporter versions and observed largely consistent splicing measurements across cell lines (Supplementary Fig. 1a-b). Hence, we selected the reporter containing *VCP* exon 10 as the upstream exon and *ACTN4* exon 16 as the downstream exon for all subsequent large-scale experiments.

To construct a representative library of exons for the screen, we took an unbiased approach. Rather than relying on exon skipping events from whole-transcriptome RNA-seq, which is noisy due to low coverage at junction-spanning reads (Supplementary Fig. 1c), we included all human internal exons ≤120-nts, along with ≥75-nts of upstream and ≥30-nts of downstream intronic sequences to preserve local splicing regulatory elements. Although introns often exceed 1 kb, most splicing regulatory elements cluster near splice sites^30^, enabling this design within DNA synthesis constraints (Supplementary Fig. 1d). To avoid nonfunctional library elements, we excluded exons that contained internal stop codons and removed sequences with low SpliceAI^14^ predicted donor/acceptor strength, as these were consistently skipped in our pilot experiments (Supplementary Fig. 1e-f). The final library comprised 46,372 native human exons spanning diverse genomic contexts, representing roughly one-quarter of all human exons (Supplementary Fig. 1g, Supplementary Table 2).

### Applying the splicing MPRA library to profile cell type-specific diversity

To ensure broad coverage of cellular diversity, we initially selected 75 cell lines from the CCLE collection spanning 10 major lineages, using Celligner transcriptomic clustering to prioritize lineage-representative groups (Supplementary Table 3)^31^. Since transfection efficiency varies substantially across cell lines^32^, we performed a pooled transfection optimization to systematically identify optimal conditions for each cell line (Supplementary Note 1). Using the optimized conditions, we transfected the pooled minigene library into each cell line individually, harvested RNA after 48 hours, and performed targeted reverse transcription followed by high-throughput sequencing (Fig. 1b, Supplementary Note 2). This allowed precise quantification of exon inclusion or skipping. All experiments were conducted in triplicate, with each sample receiving ≥20 million reads (mean: 56.0M ± 30.7M SD). On average, each sample produced 14.9 million ± 12.3 million deduplicated UMIs, corresponding to ∼320 unique UMIs per library element (Supplementary Fig. 1h). In total, we obtained high-quality splicing measurements from 43 cell lines across 10 lineages for a total of 1,993,996 splicing measurements across all cell lines (Fig. 1e, Supplementary Table 4). To quantify splicing differences, we used the metric percent spliced-in (PSI), which is the ratio of included reads divided by the sum of total skipped and included reads for each library element. Across all cell lines profiled, we observed PSI distribution to approximate a beta-distribution, where the majority of library exons have PSI close to 0 or 1, consistent with known PSI distributions (Fig. 1f). All cell lines showed strong correlation across replicates (Pearson’s r > 0.79 across triplicates, Supplementary Fig. 1i), and cell lines (Pearson’s r > 0.82 between cell lines, Supplementary Fig. 1i). There is minimal clustering between lineages, indicating that exon skipping is largely conserved across cellular contexts (Fig. 1g). Together, these results establish a robust, large-scale atlas of exon skipping across diverse cellular environments, enabling systematic discovery of regulatory principles and the design of programmable splicing elements.

### Alternative splicing shows extensive isoform complexity across cell types

Having established a large-scale atlas of exon skipping across cell types, we sought to uncover the diversity of splicing outcomes resulting from our comprehensive MPRA library. We identified a broad spectrum of alternative splicing isoforms, with a notable fraction (4.5% of all library elements) using a cryptic splice site instead of an annotated splice site as the predominant junction (Supplementary Fig. 2a). We additionally observed that a large majority of library elements (86.7%) utilized more than one splice site in at least one cell type, with 31.1% of library elements exhibited multiple alternative donor or acceptor sites in at least half of the profiled cell lines (Supplementary Fig. 2b-c). Together, these differences reflect the altered sequence context introduced by the minigene construct and highlight the advantage of a sequencing-based readout, which precisely resolves splice junctions.

Given the prevalence of cryptic splice junctions in our dataset, we aimed to investigate the sequence motifs associated with cryptic splice donors and acceptors (Supplementary Fig. 2d). Both donor and acceptor motifs closely resembled their annotated counterparts. Splice donor sites consistently featured the conserved GU dinucleotide at the +1 and +2 positions. In contrast, splice acceptor sites exhibited a strong C-and U-rich polypyrimidine tract upstream of a consensus N(CU)AG motif. These findings confirm that the cryptic and annotated splice sites identified across the library are recognized through canonical splicing signals.

### Identifying cell type-specific alternative splicing events

Next, we sought to identify library elements exhibiting cell type-specific splicing patterns. To quantify each element’s splicing specificity across cell lines, we developed a specificity score, Upsilon (υ), a scalar metric analogous to the Tau (τ) index used for tissue specificity in RNA-seq (Methods)^33^. υ ranges from 0, for library elements with uniform splicing across all cell types, to 1, for library elements with splicing restricted to a single cell type (Fig. 2a). To capture both inclusion-and exclusion-specific events – i.e., exons that are included in only one cell type and skipped elsewhere, or skipped in one cell type and included elsewhere – we additionally computed a reverse υ score by inverting PSI values. The resulting υ distributions were approximately unimodal and left-skewed, with most library elements showing low to moderate specificity, and a smaller subset exhibiting high cell type specificity, consistent with expectation that most exons are consistently regulated across diverse cellular contexts (Fig. 2b). Using υ, we identified 1004 cell type-specific library elements across the profiled cell lines, generally with cell lines representing the CNS, skin, and breast having more cell type-specific events detected than others (Fig. 2c-d, Methods).

**Figure 2:**
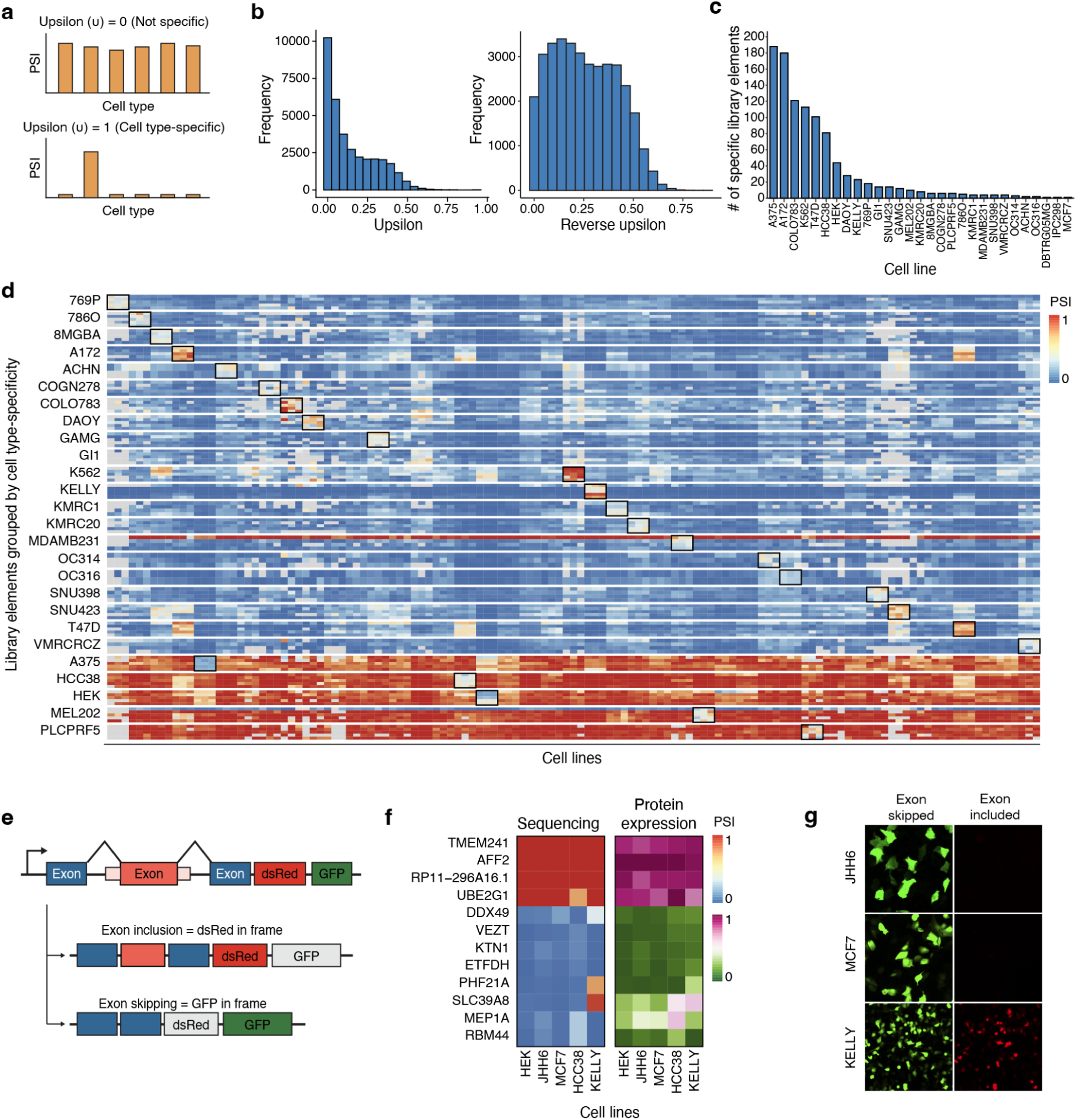
Unbiased MPRA library reveals cell type-specific splicing profiles. **a**, Description of the Upsilon metric (υ). υ = 0 corresponds to exons with ubiquitous splicing across all cell types, whereas υ = 1 corresponds to exons with highly cell type-specific splicing. **b**, Distribution of υ and reverse υ scores for all library elements. To compute reverse υ scores, all PSI values are inverted by subtracting them from 1 before υ score calculation. **c**, Distribution of the number of cell type-specific library elements identified across profiled cell lines, using υ and reverse υ score cutoffs (Methods). **d**, Heatmap of PSI values for representative cell type-specific library elements across 43 cell lines. Columns represent cell lines and rows represent library elements grouped by their cell type-specificity. Black boxes indicate the cell line in which an exon is specific. Colors denote PSI, with red indicating PSI = 1 and blue indicating PSI = 0. **e**, Frameshift reporter design for functional validation of individual library elements. Inclusion of the library exon places dsRed in frame, while exon skipping places GFP in frame. The ratio of dsRed protein expression to GFP protein expression is representative of the splicing profile in each cell line. **f**, Comparison between sequencing and protein expression readouts for constitutive and cell type-specific library elements. The left heatmap shows data from the MPRA screen, and the right heatmap shows results from protein expression frameshift reporters transfected into each cell line individually. The sequencing heatmap color scale (PSI) ranges from red (PSI = 1) to blue (PSI = 0). The protein expression heatmap color scale (normalized log_2_dsRed/GFP) ranges from pink (only dsRed expression) to green (only GFP expression). Pearson’s r = 0.933. Library elements: *TMEM241* (chr18:23399565–23399647), *AFF2* (chrX:148842965–148843002), *RP11–296A16.1* (chr15:43774475–43774518), *UBE2G1* (chr17:4331773–4331888), *DDX49* (chr19:18921662–18921748), *VEZT* (chr12:95258244–95258282), *KTN1* (chr14:55646972–55647007), *ETFDH* (chr4:158681570–158681614), *PHF21A* (chr11: 45946075–45946098), *SLC39A8* (chr4:102285948–102285973), *MEP1A* (chr6:46793551–46793576), *RBM44* (chr2:237821345-237821368). **g**, Representative images of the frameshift reporter containing *PHF21A* (chr11:45946075–45946098), expressed in JHH6, MCF7, and KELLY. Images are taken 48–72 hours post-transfection at 20x magnification.

To functionally validate whether splicing differences identified in our screen result in cell type-specific protein expression, we engineered a dual-color splicing reporter in which exon inclusion produces in-frame dsRed, while exon skipping introduces a +1 frameshift that leads to in-frame GFP expression (Fig. 2e). This design allows direct visualization of exon usage via fluorescence microscopy. We selected three groups of exons for validation: (1) four constitutively included exons, (2) four constitutively skipped exons, and (3) four cell type-specific exons. Each reporter construct was individually transfected into five representative cell lines (HEK293, JHH6, MCF7, HCC38, and KELLY), and fluorescence images were acquired after 48–72 hours. Across all three categories, reporter fluorescence closely mirrored sequencing-derived PSI values, with consistent dsRed or GFP expression patterns reflecting exon inclusion or skipping, respectively (Pearson’s r = 0.933, Spearman’s ρ = 0.796, Fig. 2f-g). Our shortlisted cell type-specific exons *PHF21A* (chr11:45946075–45946098) and *SLC39A8* (chr4:102285948–102285973) were specifically included in KELLY but skipped in all others, resulting in selective dsRed expression in KELLY alone. Conversely, constructs with *MEP1A* (chr6:46793551–46793576) and *RBM44* (chr2:237821345-237821368) drove differential inclusion and reporter expression in HCC38. These results demonstrate that the alternatively spliced library elements identified in our screen are sufficient to drive cell type-specific protein expression.

### Identifying sequence determinants driving alternative splicing through saturation mutagenesis

Alternative splicing decisions are frequently mediated by *trans*-acting RNA-binding proteins (RBPs), which recognize intronic and exonic splicing enhancers and silencers^7^. To investigate the sequence and RBP determinants underlying cell type-specific splicing, we performed single-nucleotide saturation mutagenesis on library elements specific to KELLY, a neuroblastoma cell line of the neural lineage, as KELLY-specific library elements have particularly high υ scores. We selected fifteen KELLY-specific library elements for saturation mutagenesis (Supplementary Fig. 3a) and synthesized a library spanning the entire variable region (upstream intron, skipped exon, and downstream intron) of these library elements (Supplementary Table 5). Fourteen of these elements had high PSI in KELLY and low PSI in non-KELLY cells, with one element displaying the opposite pattern. To determine the effect of each nucleotide substitution on PSI, we followed the library screening and sequencing workflow described in Fig. 1b in four cell lines: KELLY, A375, HEK293, and T47D (Fig. 3a). All cell lines exhibited strong PSI correlations across experimental replicates (n = 3, Supplementary Fig. 3b) and unmutated library elements (parent sequences) have PSI values that are highly correlated with those from the unbiased MPRA library (Pearson’s r = 0.919, Spearman’s ρ = 0.909, Fig. 3b).

**Figure 3:**
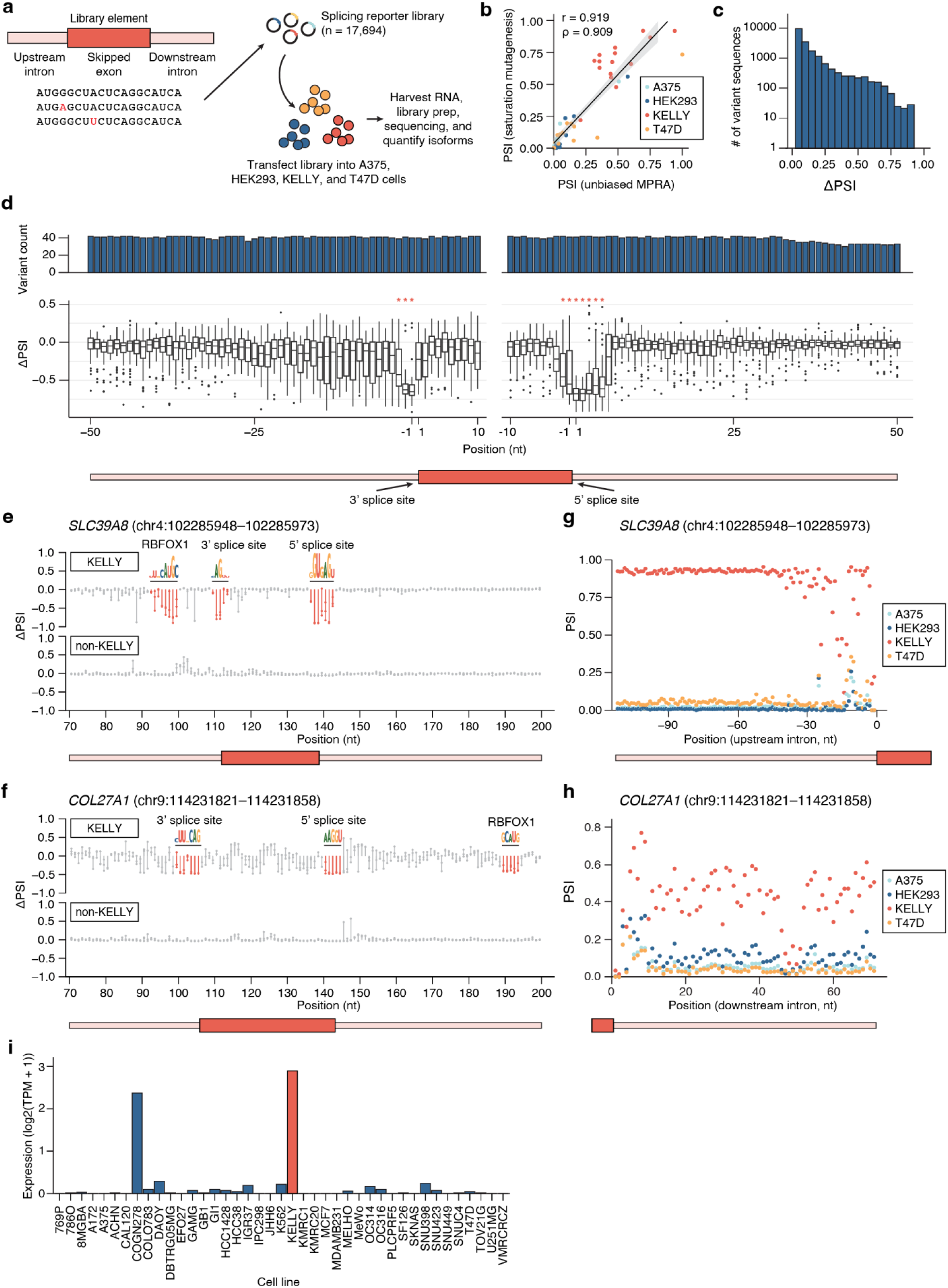
Saturation mutagenesis reveals sequence determinants driving cell type-specific splicing. **a**, Schematic of saturation mutagenesis library design and experimental setup. KELLY-specific parent sequences are selected and computationally mutagenized to create a library of single-nucleotide substitutions (n =17,694). The library is introduced into four cell lines (A375, HEK293, KELLY, and T47D), RNA is harvested, and isoform proportions are quantified by sequencing. **b**, Correlation of parent sequence PSI values between the unbiased MPRA library and the saturation mutagenesis library. Pearson’s correlation (r = 0.919) and Spearman’s correlation (ρ = 0.909) are shown; the grey shaded region indicates the 95% confidence interval. **c**, Histogram of ΔPSI between parent and variant sequences, averaged across all cell types. **d**, Position-wise ΔPSI between parent and variant sequences in KELLY (*: padj < 0.001, *z*-test per position). **e**, Effect size (ΔPSI) for all saturation mutagenesis variants in the example parent sequence *SLC39A8* (chr4:102285948–102285973), shown separately for KELLY and non-KELLY cells. Sequence motifs indicate mutation-sensitive regions identified using our *z*-score filtering approach (Methods). **f**, Effect size (ΔPSI) for all saturation mutagenesis variants in the example parent sequence *COL27A1* (chr9:114231821–114231858), shown separately for KELLY and non-KELLY cells. Sequence motifs indicate mutation-sensitive regions identified using our *z*-score filtering approach (Methods). **g**, PSI of variants across position for all four cell lines, averaged for each of the three nucleotide substitutions at each position for the upstream intron of *SLC39A8* (chr4:102285948–102285973). **h**, PSI of variants across position for all four cell lines, averaged for each of the three nucleotide substitutions at each position for the downstream intron of *COL27A1* (chr9:114231821–114231858). **i**, RNA-seq *RBFOX1* expression from the Depmap portal across target cell lines. Expression is normalized to log2(TPM + 1). TPM: transcripts per million.

To identify the individual nucleotides sensitive to mutation, we computed the difference between each variant’s PSI and its parent’s PSI to obtain an effect size (ΔPSI). Across all constructs, 80% of substitutions altered PSI by <0.1, while 2.5% had strong effects (ΔPSI > 0.5) (Fig. 3c).

High-effect substitutions predominantly localized to the annotated 5′ and 3′ splice sites, where PSI dropped to near zero upon mutation (Fig. 3d), consistent with their essential role as core spliceosomal recognition motifs. We next considered ΔPSI effect sizes across individual parent sequences, separating KELLY and non-KELLY cells. Again, we noticed that positions with large ΔPSI effect sizes clustered together and thus defined these regions with large ΔPSI effect sizes as mutation-sensitive regions using a *z*-score approach (Methods). Most of these regions overlap with splicing regulatory elements that decrease PSI, as expected (Fig. 3e-f, Supplementary Fig. 3c-o). These results confirm that the mutagenesis assay faithfully recapitulates splicing outcomes from disruptions to splicing regulatory elements.

### Mutation-sensitive regions suggest RBP binding is driving cell type-specific splicing

We were intrigued by the mutation-sensitive regions that did not cluster around the splicing regulatory elements, which we hypothesized could be RBP binding sites. To identify these potential RBPs, we compared the mutation-sensitive regions to RBP motifs from the Eukaryotic Protein-RNA Interactions (EuPRI) database^34^. We started with the exon *SLC39A8*

(chr4:102285948–102285973), the KELLY-specific parent sequence with the highest υ score (Fig. 3e,g). In KELLY, we identified three mutation-sensitive regions where mutations led to a loss in KELLY specificity. Two of these regions aligned with the annotated 3′ and 5′ splice sites, and the third region overlapped with the binding motif for RBFOX1, an RBP known to regulate alternative exon selection in neurons^35,36^. Interestingly, we found that *COL27A1* (chr9:114231821–114231858) additionally has a mutation-sensitive region that overlaps with the binding motif for RBFOX1 (Fig. 3f,h). To further explore if RBFOX1 binding may promote exon inclusion of these sequences in KELLY, we analyzed *RBFOX1* expression data from CCLE transcriptomic profiles. These results demonstrated that KELLY expressed the highest levels of *RBFOX1* among the profiled lines, with nearly all other cell types showing minimal to no expression (Fig. 3i), additionally indicating RBFOX1 may drive the selective inclusion of these two exons. Notably, COGN278 exhibited the second-highest *RBFOX1* expression despite low PSI of the *SLC39A8* exon in this context, suggesting that multiple regulatory mechanisms may work together to influence splicing outcomes.

Additionally, the saturation mutagenesis data revealed potential RBPs driving cell type-specific splicing in *SMIM8* (chr6:87330691–87330712) and *CERS6* (chr2:168766321–168766345) (Extended Figure Fig. 3i-j). Of the remaining eleven profiled parent sequences, six had mutation-sensitive regions that did not match potential RBP binders using EuPRI, and five only had mutation-sensitive regions corresponding with canonical splice site motifs. Together, saturation mutagenesis reveals both *cis-* and potential *trans-*factors important for cell type-specific splicing. These results highlight the power of our MPRA-based approach to uncover functional drivers of these sequences’ cell type specificities.

### Designing and validating a predictive machine learning model for cell type-specific splicing

Prediction of splicing outcomes would allow us to understand splicing in unmeasured cell types and sequences. While previous predictive models have been built to understand the splicing code, most have focused on predicting the location of splice sites or understanding the effects of genetic variants^13,14,37^. Furthermore, models that predict the effect of tissue-specific or cell type-specific splicing are often constrained to operate on discrete pairs of tissues, making it difficult to generalize across diverse cell types and cell states. Therefore, we set out to develop a generalized model to predict cell type-specific splicing outcomes from sequence and cellular transcriptomes.

We constructed Soma, a deep residual network^38^ that performs quantitative cell type-specific prediction for a given alternative splicing event using both sequence and gene expression information. Soma was designed to address two key tasks: (i) predicting PSI for unseen sequences and (ii) predicting PSI in unseen cell types (Fig. 4a, Supplementary Fig. 4a). To represent diverse cell types and cell states, we used gene expression profiles as continuous descriptors of cellular identity. We took inspiration from modelling approaches used in dissecting CREs to construct our model (Fig. 4b)^39^. First, we performed pretraining of a convolutional neural network (CNN) using only sequence information to predict the mean PSI of an unseen sequence across all cell types. Our training was performed using inputs of 250-nt splicing reporter sequences and outputs of PSI, which captures the cell type-agnostic effects of splicing. To perform fine-tuning of the model, we input both sequence and cell type gene expression information (log-normalized expression counts), where we trained the model on the residuals between the mean and individual cell types (Methods). In Soma’s first task of predicting PSI for unseen sequences, we observed that the prediction accuracy solely using sequence information alone resulted in a Pearson’s correlation between predicted and measured PSIs of Pearson’s r = 0.695 (Supplementary Fig. 4b). When we incorporated the gene expression information, the prediction accuracy increased to Pearson’s r = 0.730 (Fig. 4c). This result was consistent across train-test splits (Fig. 4d, Methods). For each test sequence, predicted and observed PSI values across cell lines were closely aligned (Fig. 4e). Together, these results indicate that Soma is learning the impact of RBP expression on cell type-specific splicing regulation.

**Figure 4:**
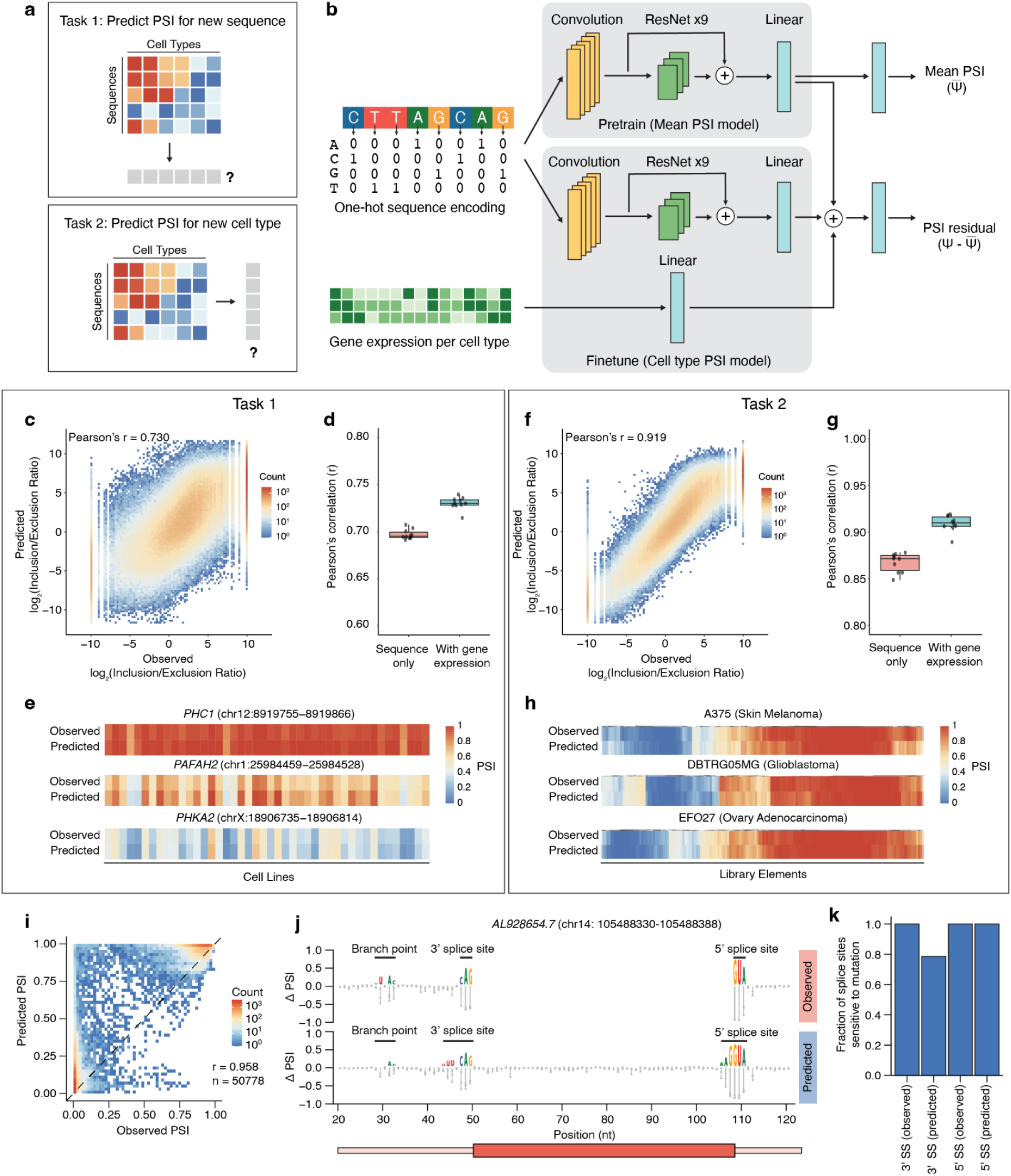
Soma accurately predicts splicing outcomes across unseen sequences and cell types and identifies mutation-sensitive motifs. **a**, Overview of Soma tasks. Task 1 involves predicting PSI for unseen sequences, and Task 2 involves predicting PSI for unseen cell types. **b,** Schematic of the Soma architecture. A convolutional residual network is pretrained to predict mean PSI across cell types and subsequently fine-tuned with gene expression inputs to predict cell type-specific PSI. **c**, Correlation between predicted and observed splicing outcomes for held-out sequences. Each point represents a single held-out sequence. **d,** Pearson’s correlations across ten train-test splits comparing sequence-only and sequence with gene expression models when predicting unseen sequences. **e,** Representative examples of observed versus predicted PSI across cell types for individual sequences. **f**, Correlation between predicted and observed splicing outcomes for held-out cell types. Each point represents a single held-out sequence. **g**, Pearson’s correlations across ten train-test splits comparing sequence-only and sequence with gene expression models when predicting unseen cell types. **h,** Representative examples of observed versus predicted PSI across sequences for individual cell types. **i**, Predicted versus observed PSI values for single-nucleotide saturation mutagenesis libraries. The constitutively spliced library was assayed in seven cell lines (8MGBA, HCC38, JHH6, KELLY, KMRC20, MCF7, T47D). Pearson’s correlation r = 0.958. Spearman’s correlation ρ = 0.821. **j**, Effect size ΔPSI for all variants in example parent sequence AL928654.7 (chr14: 105488330–105488388), split between observed and predicted data. PSI values are averaged across the seven tested cell lines. Sequence motifs are mutation-sensitive regions called by the *z*-score filtering approach. **k**, Fraction of the eleven parent sequences that correctly identify splice sites as splicing sensitive regions, in both the measured and predicted datasets.

For the second task, we asked Soma to predict the splicing outcome of a sequence in an unseen cell type. We observed a high correlation (Pearson’s r = 0.866) across unseen cell types (Supplementary Fig. 4c), which again increased (Pearson’s r = 0.919) when Soma incorporated cell type expression data (Fig. 4f). This result was also consistent across 10 test-train splits (Fig. 4g) and similar to task 1, predicted and observed PSI values across sequences were closely aligned (Fig. 4h). This demonstrates Soma’s ability to use gene expression data to extend splicing predictions across non-discrete cell type categories.

We next considered whether the model can identify splicing regulatory elements. To assess this, we designed another saturation mutagenesis library focused on 23 constitutively-spliced parent sequences (eleven with high PSI and twelve with low PSI) and assayed splicing in seven representative cell lines (Supplementary Table 6). Concurrently, we asked the model to predict the splicing response to each of these mutated sequences in each cell line. Overall, the correlation between the predicted and observed PSI levels was high (Pearson’s r = 0.958, Spearman’s ρ = 0.821, Fig. 4i). To find splicing regulatory elements affected by mutagenesis in both the predicted and observed data for the high PSI sequences, we performed *z*-score filtering to identify mutation-sensitive regions (Methods). The model correctly predicted mutation-sensitive regions within the profiled sequences, including the 3′ splice site, 5′ splice site, branch point, and polypyrimidine tract (Fig. 4j). Across the profiled parent sequences with high PSI, the vast majority of 3′ and 5′ splice sites were identified as mutation-sensitive regions in both the predicted and observed data (Fig. 4k). Together, these results indicate that the model can both accurately predict splicing outcomes and understand core splicing sequence grammar.

### Designing and validating a generative AI model for cell type-specific splicing

The ability of our Soma model to predict cell type-specific splicing patterns enables its use as a “teacher” that can be used to train generative deep learning models that can design new RNA splicing sequences with specific splicing outcomes or cell type-specificity. To do so, we constructed Melange, a generative AI model that can generate new sequences with desired splicing outcomes. We used this model to perform two tasks: (i) generate sequences with user-defined PSI values and (ii) generate sequences with cell type-or cell state-specific activity (Fig. 5a).

**Figure 5:**
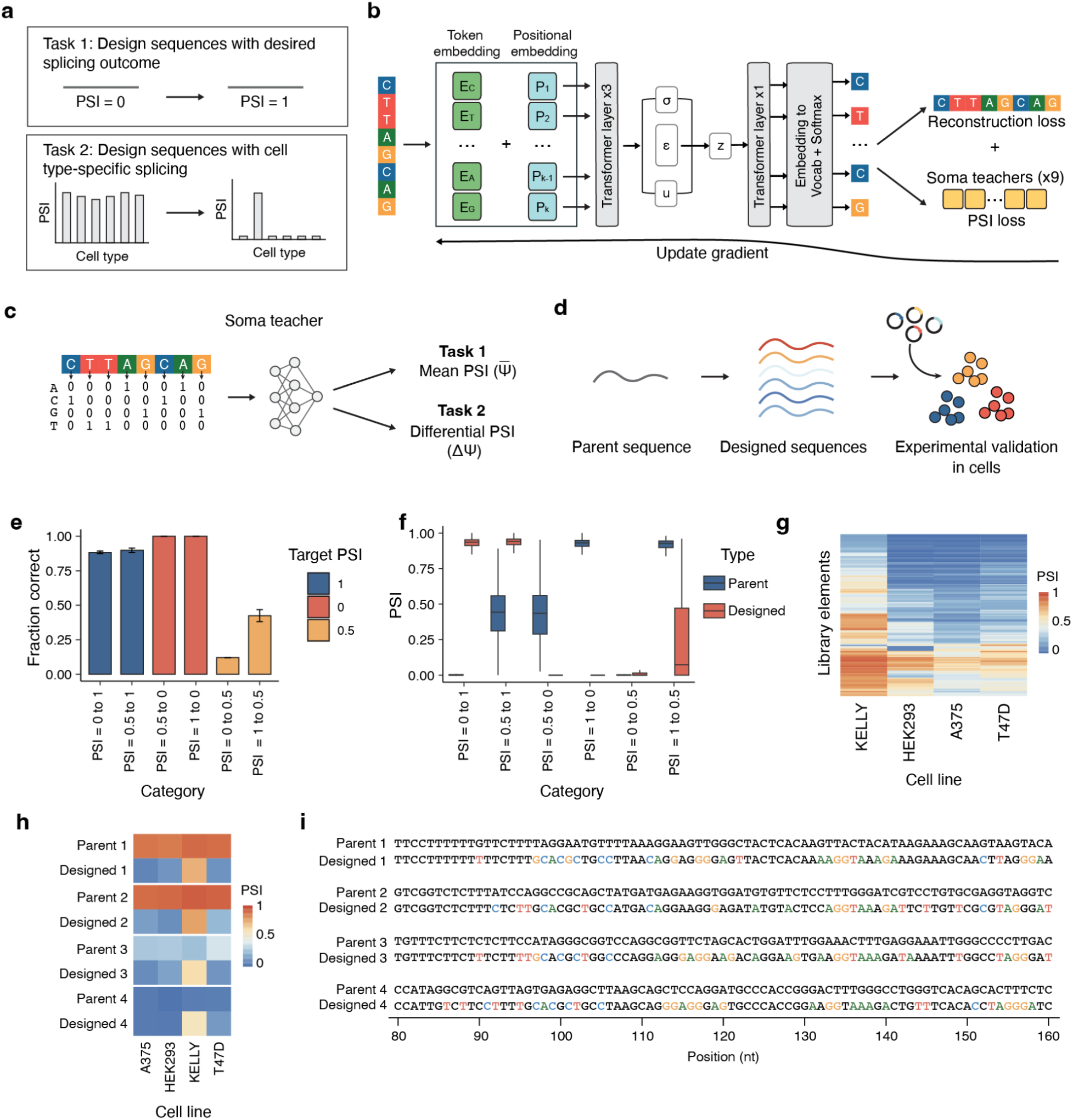
Melange generates synthetic splicing sequences with programmable cell type-specific activity. **a**, Schematic of design tasks given to the Melange model. Task 1 involves designing constitutively spliced sequences to achieve user-defined PSI values, while Task 2 involves designing sequences with cell type-specific splicing. **b**, Melange architecture. A transformer-variational autoencoder (VAE) integrates guidance from nine independently trained Soma teacher models to generate sequences with desired splicing outcomes. One held-out Soma teacher is used for filtering designed sequences. **c,** Teacher-student framework. Soma teachers constrain Melange during sequence generation to ensure quantitative and cell type-specific control of splicing. **d**, Workflow for experimental validation. Designed libraries were synthesized and tested in A375, HEK293, KELLY, and T47D. **e**, Fraction of designed sequences achieving the intended outcome across design categories in HEK293. Error bars represent the standard deviation across experimental replicates (n = 3). **f**, Distribution of PSI values for designed sequences stratified by target category in HEK293, showing both the parent and designed sequences. **g**, PSI values of all KELLY-specific elements generated by Melange across profiled cell lines, filtered for significant events identified by rMATS. **h**, PSI values of representative KELLY-specific elements comparing parent and designed variants. **i**, Example sequences from (h). Highlighted nucleotides indicate substitutions between the parent and designed sequences.

We trained Melange by combining a transformer-based variational autoencoder (VAE)^40,41^ with guidance from Soma “teacher” models (Fig. 5b-c). This design allows Melange to generate novel variants that introduce mutations at informative positions while maintaining overall sequence integrity. To enhance robustness, we adopted an ensemble learning strategy, training ten independent Soma teachers with distinct data partitions and random initializations. Nine models were aggregated during training to capture complementary decision boundaries, while one model was held out as a filtering module to ensure that only high-confidence designs progressed to downstream testing. For both user-defined PSI values and cell type-specific design tasks, we validated Melange-generated sequences experimentally across four representative cell types: A375, HEK293, KELLY, and T47D (Fig. 5d).

Our first task was to generate sequences with a user-defined constitutive PSI across several cell types. To test this, we stratified sequences into three bins: low (PSI < 0.2), intermediate (0.2–0.8), and high (>0.8). The model was trained on these bins and tasked with converting seed sequences between categories. For example, 200 low-PSI sequences were used as inputs to generate high-or intermediate-PSI variants. We synthesized a validation library and assayed splicing in the four cell types (Supplementary Table 7). We observed that across bins, more than 90% of generated sequences achieved the intended outcome, with lower accuracy for conversions to the intermediate bin, likely reflecting higher biological variance and less training data within this range (Fig. 5e, Supplementary Fig. 5a-b). Among sequences that were designed to either low or high bins, the PSI values of tested sequences clustered tightly around the designated range (Fig. 5f). Each designed sequence exhibited consistent PSI across the measured cell lines and contained numerous substitutions relative to its parent sequence (Supplementary Fig. 5c-d). Our results show that Melange can design sequences across a range of designated PSI levels.

In the second task, we asked Melange to design sequences with selective activity in a single cell type. To achieve this, we trained the model on PSI differences between KELLY and non-KELLY cells using our unbiased MPRA library. We then asked the model to generate KELLY-specific sequences from a set of parent sequences that are not KELLY-specific (Supplementary Table 8). We experimentally validated these designed sequences (n = 2,000) in four cell types (A375, HEK293, KELLY, and T47D), and identified 133 sequences with significant KELLY specificity (Fig. 5g-i, Supplementary Fig. 5e). These sequences accounted for 6.7% of the designed set, representing a 33-fold enrichment compared to the baseline frequency of KELLY-specific sequences in the unbiased MPRA screen (0.2%). Most of the designed sequences exhibited higher PSI in KELLY than in non-KELLY cells, and many showed PSI changes equal to or greater than the strongest endogenous KELLY-specific exons. Together, these results demonstrate that Melange can generate *de novo* cell type-specific sequences, expanding the design space for cell type-specific splicing control.

### Iterative design of sequences with programmable splicing in splicing factor-mutated cells

To demonstrate the power of SPICE as an integrated framework for large-scale splicing profiling and generative design, we applied it to design sequences specific to cancer-associated splicing factor mutations (Fig. 6a). Such mutations, including hotspots in *SF3B1* and *U2AF1* and loss-of-function events in *FUBP1*, *RBM5*, *RBM10*, and *ZRSR2*, are recurrent in myelodysplastic syndromes, acute myeloid leukemia, chronic myelomonocytic leukemia, and multiple solid tumors (Fig. 6b)^8,42–45^. Mutation or dysregulation of these splicing factors results in patterns of missplicing in hundreds of genes and can be leveraged to engineer selective targeting of gene therapies or toxic payloads specifically to splicing factor mutant cancer cells.

**Figure 6:**
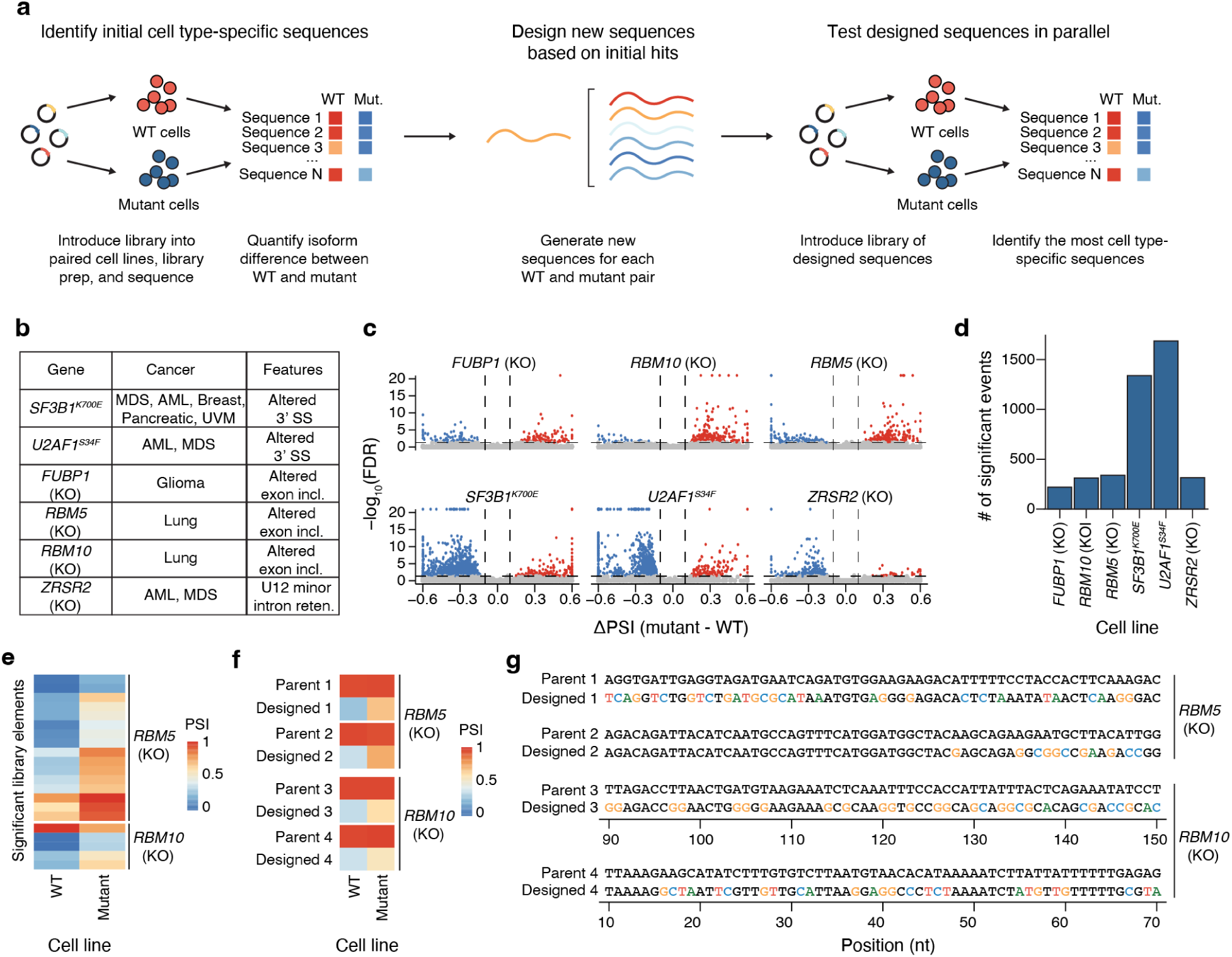
SPICE enables the design of cancer mutation-specific splicing elements. **a**, Schematic of the integrated experimental–computational framework for designing splicing sequences specific to cells harboring splicing factor mutations. Cell lines carrying defined splicing factor mutations were first generated, and a pooled minigene library was introduced into paired wild-type (WT) and mutant cell lines. RNA was harvested, and isoform proportions were quantified by sequencing. These data were used to train the Melange model, which subsequently designed new cell type-specific sequences. The designed sequences were synthesized, reintroduced into the same WT and mutant cell lines, and experimentally validated to identify mutation-specific splicing elements. **b**, Splicing factor mutations profiled in this study, their associated cancer types, and characteristic splicing alterations. **c**, Volcano plots of significantly differentially spliced library elements (ΔPSI) for each mutant versus wild-type cell line. Significant events have ΔPSI > 0.1 and FDR < 0.05. Red points indicate higher PSI in knockout (KO) cells than in WT cells, and blue points indicate lower PSI in KO cells. **d**, Number of significant library elements identified per mutant background, filtered by ΔPSI > 0.1 and FDR < 0.05. **e**, Heatmap of PSI values for significant designed sequences specific to *RBM5* and *RBM10* KO cells (ΔPSI > 0.1, FDR < 0.05). **f**, Heatmap showing PSI values for representative parent-design pairs across WT and mutant cell lines. **g**, Representative examples of paired parent and designed sequences. Nucleotides altered relative to the parent sequence are highlighted.

Using our unbiased MPRA library, we profiled exon skipping across matched wild-type and engineered mutant cells carrying splicing factor mutations. We evaluated 6 splicing factor mutations commonly observed in cancer, including hotspot mutations in *SF3B1*^K700E^, *U2AF1*^S34F^, and CRISPR-engineered loss of function in *FUBP1*, *RBM5*, *RBM10*, and *ZRSR2* (Fig. 6b, Supplementary Table 9). We identified dozens to hundreds of library elements that differed in PSI (Fig. 6c-d) or alternative splice site selection (Supplementary Fig. 6d-e) between the wild type and mutant cell lines and recovered patterns of mis-splicing consistent with known splicing factor mutant phenotypes. For the loss of function alterations, we found 311 and 339 differentially spliced elements with a predominance in increased exon inclusion in *RBM5* and *RBM10* KO cells, respectively (Fig. 6d). *RBM5* and *RBM10* are paralogous genes that promote skipping of cassette exons, particularly in introns with cytosine predominant polypyrimidine tracts^46^. These results confirm that our MPRA approach can recover known patterns of mis-splicing in splicing factor-mutant contexts.

Next, we tasked Melange with designing *de novo* sequences with differential splicing between mutant and wild-type contexts. We chose to focus on *RBM5* and *RBM10* loss-of-function mutation contexts, as the features that drive mutant-associated mis-splicing are less well explored compared to other splicing factor mutations. We seeded Melange with 581 constitutively spliced parent sequences and asked the model to generate either *RBM5*-specific or *RBM10*-specific sequences. We experimentally validated these designed sequences in both wild-type and mutant cell lines and subsequently recovered 1,605 *RBM5*-and 415 *RBM10*-specific sequences with high-quality sequencing reads (Supplementary Table 10). We identified sixteen *RBM5*-specific and five *RBM10*-specific sequences, both representing about a 5-fold enrichment in the frequency of specific sequences compared to baseline (1% for *RBM5*, 1.2% for *RBM10*, and 0.2% for the unbiased MPRA library, Fig. 6e-g, Supplementary Fig. 6f). The significant designed sequences have PSI values ranging from zero to one, though almost all tend to have higher PSI in the mutant cell line than in the WT cell line. This result is concordant with prior literature showing the role of RBM5 and RBM10 in promoting exon skipping^46^.

Together, our results demonstrate the integrated capability of SPICE to profile, predict, and generate splicing elements. This framework enables the discovery of cell type-specific sequences and the design of synthetic elements with programmable activity in defined cellular states.

## Discussion

Here we introduce SPICE, an integrated framework for the systematic profiling and generative design of sequences governing cell type-specific alternative splicing. By performing an MPRA across 43 diverse cell lines and six engineered splicing factor mutant lines, we generated the largest functional datasets of exon skipping to date. From this atlas, we identified hundreds of sequences with cell type-specific activity and uncovered sequence features that underlie their specificity. Leveraging these data, we trained a predictive model, Soma, that accurately predicts splicing outcomes across cellular contexts and a generative model, Melange, that designs synthetic elements with programmable splicing behavior. Large-scale experimental validation confirmed the robustness of these designed sequences, positioning SPICE as both a resource for dissecting splicing biology and a framework for engineering gene regulation.

A unique feature of SPICE is its capacity to generate novel splicing sequences, establishing it as the first framework to integrate large-scale experimental data generation with the *de novo* design of cell type-specific splicing elements. While prior models of splicing have focused on predicting splice site choice or variant effects, our generative model Melange demonstrates that splicing regulation can also be programmed in a cell type-specific manner. By conditioning sequences designed on Soma-derived constraints, Melange achieves quantitative control of exon inclusion and generates elements with cell-type-or mutation-specific activity at rates far exceeding random genomic discovery. This represents a shift from interpreting splicing regulation to actively engineering it. Additionally, unlike approaches that treat cell types as discrete categories, our framework represents cellular identity through gene expression profiles, better reflecting the continuum of cellular states.

SPICE also has several limitations. First, our experiments were conducted in established cancer cell lines, which may not fully recapitulate primary or *in vivo* contexts. Future studies will need to extend into diverse primary cell types and physiological settings. Second, although our MPRA library was relatively large at 46,372 elements, it still sampled only a small fraction of the possible sequence design space. By contrast, existing models for CRE design were trained on datasets exceeding 770 million short MPRA sequences^4^. Access to similarly large-scale datasets for splicing would enable more comprehensive mapping of regulatory grammars and support the development of models with stronger predictive and generative capabilities.

Looking ahead, SPICE could be extended to other modes of alternative splicing, such as alternative 5′ or 3′ splice sites, intron retention, and mutually exclusive exons. It is also well-positioned to target disease-associated cell states that remain difficult to access with conventional approaches, particularly in immunology, cancer, and rare disease contexts. Our framework could also be integrated with other mechanisms of cell type-specific control to achieve precise, combinatorial targeting^1–5^. The short, modular elements identified and generated by SPICE are additionally compatible with viral vectors and other delivery platforms, making them directly translatable into therapeutic or reporter constructs. By uniting large-scale profiling, predictive modeling, and generative design, SPICE provides both a rich dataset for probing the biology of splicing and a versatile framework for constructing compact, programmable elements. We anticipate that future integration with larger datasets, additional splicing modalities, and *in vivo* delivery systems will expand the scope of programmable splicing control, positioning SPICE as a foundational tool for precision gene regulation and therapeutic engineering.

## Materials and methods

### Mammalian cell culture

HEK293FT cells (referenced as HEK293 in the text and figures, Thermo Fisher - R70007) and A375 cells (ATCC) were cultured in Dulbecco’s Modified Eagle Medium with GlutaMAX (Thermo Fisher Scientific - 10564011) supplemented with 10% (v/v) fetal bovine serum (FBS, Sigma-Aldrich F4135) and 1x penicillin-streptomycin (Thermo Fisher Scientific 15140122).

K562 cells (ATCC, CCL-243) were cultured in RPMI 1640 medium with GlutaMAX (Thermo Fisher-61870036) supplemented with 10% (v/v) FBS and 1× penicillin-streptomycin. All other cell lines were obtained from the PRISM Platform at the Broad Institute (Supplementary Table 3). The PRISM cells were cultured in the same supplemented RPMI 1640 medium. Adherent cells were maintained at confluency below 80%–90% at 37°C and 5% CO_2_. Suspension cells were maintained at confluency below 1.5 × 10^6^ cells/mL at 37°C and 5% CO_2_.

### Design of splicing MPRA minigene reporter

To determine the optimal reporter design, we first tested three reporter versions that differed in their 5′ splice site (5′ss) strength, as flanking splice site strength can modulate the effect size in minigene reporters^28^. We quantified 5′ss strength using MaxEntScan, a computational model that scores splice sites based on maximum entropy principles of sequence motifs^47^. Specifically, we used upstream exons with strong (*VCP* exon 15, MaxEntScan = 10.15), intermediate (*EMC7* exon 3, MaxEnt = 8.35), and weak (*VCP* exon 10, MaxEntScan = 6.66) 5′ splice sites. All reporters had the same *ACTN4* exon 16 as the downstream exon. We first performed a pilot screen with a small minigene library of 120 test exons transfected into three cell lines. RNA was harvested 48 hours post-transfection, followed by targeted library preparation and sequencing (Supplementary Table 1). To quantify splicing differences, we use the metric percent spliced-in (PSI), which is the ratio of included reads divided by the sum of total skipped and included reads for each library element. Since we observed largely consistent splicing measurements across cell lines, we decided to randomly select the weak 5′ss reporter for our full screen.

### Design of unbiased splicing MPRA library

Genomic exon annotations were obtained from GRCh38 (Gencode v39). Unique exons were filtered to retain sequences 10–120 nucleotides in length that lacked in-frame stop codons and contained BsmBI sites compatible with cloning. SpliceAI was applied to each exon with its flanking minigene context to predict splice donor and acceptor probabilities. To determine an appropriate prediction threshold, a logistic regression model was trained on a pilot set of 120 exons, categorized as either “always skipped” (PSI < 0.05) or “always included” (PSI ≥ 0.95). Exons predicted to be constitutively skipped were excluded, yielding a final library of 46,382 elements. For exons that passed filtering, flanking intronic regions were trimmed to ensure a minimum length of 75-nt of upstream intron and 30-nt of downstream intron were included, for a total sequence length 215-nt per element. Each library element was assigned a 14-nt barcode randomly selected from a validated set ^48^.

### Cloning of splicing libraries

Library elements were purchased as an oligonucleotide pool (Twist Biosciences) and amplified using KAPA HiFi ReadyMix Hot Start (Roche - KK2601). Cloning of the libraries happened using two step-wise cloning steps. First, the oligonucleotide pool was cloned into a custom-designed parent plasmid vector that contained the constant upstream exon and intron sequences (pDC297, Supplementary Table 1) using NEBuilder HiFi DNA Assembly Master Mix (NEB - E2621L). The library was transformed into Endura Electrocompetent Cells (Sigma-Aldrich - LGC6024221) and plated onto five 30-cm carbenicillin-containing agar plates to grow overnight. The next day, cells were scraped off and the plasmid DNA was harvested using the ZymoPURE II Plasmid Maxiprep Kit (D4203). To ensure sufficient library diversity, the total number of colonies was estimated using serial dilutions of the transformed bacteria.

After the first transformation, a second cloning step was performed to add in the constant downstream intron and exon sequence using a gBlock (IDT; DCG197) and the NEBridge Golden Gate Assembly Kit (BsmBI-v2, E1602L). The same transformation protocol was used again.

After each of the steps of library amplification and maxiprep, the plasmid pool was sequenced on the MiSeq (Illumina) to ensure sufficient sequence diversity.

### Transfection and electroporation of splicing libraries into cell lines

The day before transfection, adherent cells were seeded into 6-well plates (Corning) or 10-cm plates (KMRC1 cells, Corning) at a density so that the cells are approximately 70% confluent 24 hours later. Cells were transfected using TransIT-LT1, TransIT-X2, or TransIT-2020 (Mirus Bio; Supplementary Table 3), following the manufacturer’s instructions. Each well was transfected with 2.5 μg of splicing library plasmid, except for KMRC1 cells, which received 15 μg, with three biological replicates per condition. Suspension cells were electroporated using the P3 Primary Cell 4D-Nucleofector kit (Lonza, V4XP-3032). For each electroporation, 2 × 10^6^ cells were transfected with 4 μg of splicing library plasmid. Following transfection or electroporation, cells were cultured for 48 h prior to RNA extraction. Optimal transfection and electroporation parameters were established in pilot experiments before large-scale library introduction (Supplementary Note 1).

### Extraction of RNA and next-generation sequencing of amplicon splicing libraries

Total RNA was extracted from transfected or electroporated cells using the RNeasy Plus Mini Kit (Qiagen, 74134), with on-column DNase digestion (Qiagen, 79254) to remove residual plasmid and genomic DNA. cDNA synthesis was performed with Maxima H Minus reverse transcriptase (Thermo Fisher, EP0743) using a library-specific primer containing a 10-nt unique molecular identifier (UMI). cDNA was purified using 1.5x AMPure SPRI beads (Beckman Coulter). To avoid overamplification, the number of PCR cycles was determined by qPCR of cDNA (Luna Universal qPCR, NEB, M3003).

Libraries were generated using either a two-step or one-step PCR strategy. In the two-step protocol, cDNA was first amplified with Q5 Hot Start High-Fidelity 2X Master Mix (NEB M0494) with the following cycling conditions: 95°C for 30 s; predetermined number of cycles (Ct + 3 cycles) of 98°C for 15 s, 72°C for 15 s, and 72°C for 1 min; final extension at 72°C for 2 min. PCR products were purified with 1.5x SPRI beads, then amplified with sample index primers for 12 cycles using Q5 Hot Start High-Fidelity 2X Master Mix using the same PCR protocol. In the one-step protocol, cDNA was amplified directly with Titanium Taq (Takara, 639210) using a mixture of one-step PCR P5 primers and a one-step PCR P7 index primer, under the following cycling conditions: 95°C for 5 min; predetermined number of cycles of 95°C for 30 s, 66°C for 30 s, and 72°C for 1 min; final extension at 72°C for 5 min. PCR products were purified with a double-sided AMPure SPRI cleanup (0.5× reverse elution followed by 1.2× forward elution). Primer sequences used can be found in Supplementary Table 1.

Libraries were quantified with the Qubit 1X dsDNA High Sensitivity Assay Kits (Thermo Q32851) and checked for purity with Tapestation using the high sensitivity D5000 kit (Agilent - 5067-5592). Amplicons were pooled and prepared for sequencing on either a Novaseq X or NextSeq 1000 (Illumina) with paired-end reads (read1, 217bp; index1, 8bp; index2, 8 bp; read2, 60bp). 20% PhiX was added to the sequenced libraries and approximately 1000x coverage was given to each cell line and library exon. Reads were demultiplexed and analyzed with appropriate pipelines.

### Processing of amplicon splicing library NGS reads

Raw paired-end FASTQ files were processed to quantify the exon inclusion levels for each reporter element. For each read, R1 contained the reporter sequence spanning the exon-exon junctions, while R2 contained the barcode for each element. The barcode and UMI from each sequence was first extracted using UMI-tools (v1.1.2) using a whitelist of known barcodes.

Reads not matching the barcode whitelist were discarded. A custom Python script was used to map each reporter sequence to its corresponding splicing outcome. UMI-level counts were deduplicated and aggregated to produce element-barcode count matrices. To account for batch effects from barcode swapping, we implemented a correction for chimeric reads (Supplementary Note 2). This correction was applied to all libraries prior to downstream quantification of exon inclusion.

### Identification of cell type-specific elements

We quantified cell type specificity using two complementary metrics, Tau (τ) and Upsilon (υ). Tau is a widely used measure of tissue specificity in RNA-seq data, where read counts are approximately normally distributed. However, in splicing assays, where PSI values for splicing follow a beta distribution, Tau can overestimate specificity for elements with limited dynamic range. To address this, we developed a second metric, Upsilon, which is more robust to differences in variance and dynamic range across cell types. For each library element, Tau and Upsilon scores were calculated on PSI values across all cell types, ranging from 0 (not cell type specific) to 1 (exclusive to a single cell type). The formulas were as follows:

The index υ was defined as:

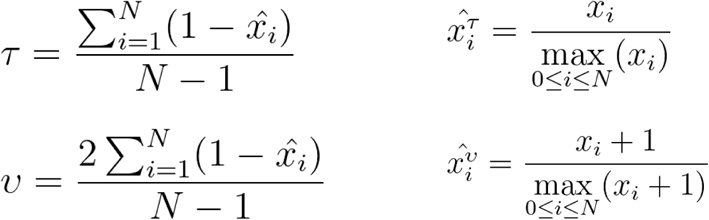

where *N* is the number of cells and 𝑥𝑖 is the expression profile component normalized by the maximal component value.

We defined cell type–specific elements based on conservative cutoffs for both Tau and Upsilon, ensuring robustness to spurious calls from low dynamic range. Specifically, elements were shortlisted if they satisfied any one of the following conditions:

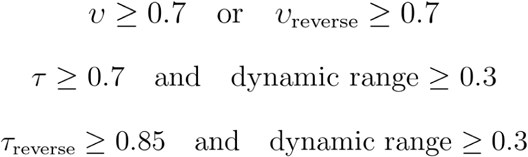

where dynamic range is defined as the difference between the maximum and minimum PSI for each cell line. Shortlisted elements were ranked by decreasing Upsilon score to prioritize those with the strongest evidence of cell type-specific regulation.

### Splicing analysis of CCLE RNA-seq data

Publicly available transcriptomic RNA-seq data from CCLE cell lines^49^ were downloaded from the Terra CCLE-public workspace (https://app.terra.bio/#workspaces/broad-firecloud-ccle/CCLE-public). Raw reads were extracted from the BAM files and aligned to GRCh38 using HISAT2^50^. Differential analysis of splicing events was identified using rMATS^51^.

### Validation of individual library elements using fluorescence reporters

Frameshift reporters of individual library elements were generated by Gibson assembly of synthetic minigene sequences (IDT) into a modified pSLED backbone. The parent pSLED.NPL was a gift from Jonathan Ling (Addgene plasmid #193014; http://n2t.net/addgene:193014; RRID: Addgene_193014). This reporter was developed to display different ratios of dsRed and GFP based on PSI. The library element was placed upstream of both fluorescent genes. Two versions of the frameshift reporter were designed, based on the exon length within the library element. If the library element contained an exon that would lead to a +1 frameshift, exon inclusion would lead to expression of dsRed and exon skipping would lead to expression of GFP.

For exons that led to a +2 frameshift, the design was switched, such that exon inclusion leads to expression of GFP and exon skipping leads to expression of dsRed. Downstream of the dsRed and GFP sequences is the IRES driving BFP expression, used as a transfection normalization control.

Validation of individual elements was performed by transfecting the reporter into different cell types, then imaging the cells at 48 hours post-transfection on a Nikon Ti2-E confocal microscope at room temperature. Images were collected at 10× magnification with fixed acquisition settings, and representative images were also obtained at 20×. Images were collected in 405 nm (50% power, 100 ms exposure), 488 nm (50% power, 100 ms exposure), and 561 nm (50% power, 500 ms exposure). BFP signal was used for focusing and normalization.

Microscopy images were processed in MATLAB using watershed segmentation to quantify per-cell fluorescence of BFP, dsRed, and GFP. For +1 frameshift reporters, GFP:dsRed ratios were used as proxies for PSI, and the inverse ratio for +2 reporters. Ratios were normalized against control elements with defined PSI values (0 or 1), transformed to a logarithmic scale, and finally rescaled between zero and one for downstream analysis.

### Design of KELLY-specific saturation mutagenesis library

The KELLY-specific saturation mutagenesis library (Supplementary Table 5) was derived from fifteen parent sequences identified as the most highly KELLY-specific. Each library element was computationally mutagenized to include all possible single-nucleotide substitutions, spanning the entire upstream intron, exon, and downstream intron regions. This design yielded a total of 9,669 unique variants, ensuring full coverage of single-base changes at every position. Each library element was assigned a 14-nt barcode randomly selected from a previously designed list^48^. The library was cloned using the same steps as described above.

### Identification of mutation-sensitive regions and potential relevant RBPs from saturation mutagenesis data

Enriched sequence regions were identified in the saturation mutagenesis data using a z-score filtering approach. First, we separated out variants by parent sequence and cell line, then identified the effect size (ΔPSI = parent PSI - variant PSI) for each mutated sequence, and found the mean and standard deviation of effect sizes for each set. Next, we smoothed our data using a Gaussian filter and obtained the baseline sensitivity of each parent sequence to mutations by computing the 25–75% IQR. We labeled positions with a z-score of at least three (KELLY-specific library) or four (constitutive library) away from the median as mutation-sensitive positions. From these individual positions, we then grouped them into blocks of a minimum size of three (KELLY-specific library) or two (constitutive library) and with a minimum mean effect size of ΔPSI ≥ 0.25 (KELLY-specific library) or ΔPSI ≥ 0.25 (constitutive library), which we label as mutation-sensitive regions.

### Identification of associated RBP motifs

To identify 3′ and 5′ splice sites in the mutation-sensitive regions identified previously, all mutated parent sequences were first mapped back to annotated splice junctions. To find potential RBPs driving the splicing of these parent sequences, any mutation-sensitive regions remaining for each parent sequence were next imported into the Eukaryotic Protein-RNA Interactions (EuPRI) database ^34^ using the CISBP-RNA webserver. We used the RNA Scan tool to identify RBP binding motifs, filtering the results to human RBPs with PWM LogOdds scores of at least 10.

### Design of constitutively-spliced saturation mutagenesis library

The constitutively-spliced saturation mutagenesis library (Supplementary Table 6) was derived from 23 constitutively-spliced library elements identified from the unbiased MPRA library, with eleven having PSI values close to one, and twelve having PSI values close to zero. To allow for more parent sequences to be screened, only up to 60-nts upstream of the 3′ splice site and up to 40-nts downstream of the 5′ splice site were mutagenized with single-nucleotide saturation mutagenesis. Only exons with 35 or fewer nucleotides were included in the mutagenesis. This design yielded a total of 5,810 unique variants. Each library element was assigned a 14-nt barcode randomly selected from a previously designed list^48^. The library was cloned using the same steps as described above.

### Dataset Preprocessing for the development of machine learning models

Raw experimental counts were converted into log-ratios of exon inclusion versus exclusion. For each sequence, counts were normalized to a total of 1,000 reads, and the log-ratio was calculated as: log_2_((I+1)/(E+1)), where I and E represent inclusion and exclusion counts, respectively.

Log-ratio was used instead of PSI because it approximates a normal distribution, facilitating downstream modeling. Values were averaged across cell lines and mapped to the corresponding sequence barcodes. DNA sequences were then one-hot encoded, representing each nucleotide (A, T, C, G) as a four-dimensional binary vector. To ensure consistent input length, sequences were padded with the constant reporter backbone to 250-nts.

### Computational Environment for modelling

All models were implemented using PyTorch and trained on a server with an AMD Ryzen Threadripper PRO 3975WX CPU and an NVIDIA RTX A6000 GPU. PyTorch utilized CUDA libraries for GPU acceleration. Software dependencies included Python 3.9, PyTorch 1.12, NumPy, Pandas, and other standard Python scientific computing libraries. Random seeds were fixed throughout all training and evaluation processes to ensure reproducibility of results.

### Development of the Soma Model

A convolutional neural network (CNN) was designed using an architecture with residual connections to predict continuous PSI values from DNA sequences. The model consisted of multiple convolutional layers organized into three sequential residual blocks. Each block contained multi-scale convolutional kernels of sizes 5, 11, and 21 to capture nucleotide sequence patterns at different resolutions. Batch normalization and dropout (rate = 0.2) were applied after each convolution to enhance generalization. Residual connections within each block facilitated gradient flow and improved training stability. After passing through these convolutional layers, outputs were aggregated through fully connected layers to yield final PSI predictions as continuous values.

To further integrate gene expression features for predicting cell type-specific PSI, we incorporated an additional fully connected neural network to encode gene expression profiles. The resulting gene latent representations were concatenated with the sequence-derived latent vectors from the CNN, enabling joint modeling of sequence patterns and gene-level regulatory context. This fused representation was then passed through additional fully connected layers to predict the deviation of PSI values for each cell type relative to the average PSI across cell types.

For model training, the PSI matrix (sequences × cell types) was partitioned along two axes according to the training task. In the sequence split, 80% of sequences were used for training and 20% were held out for testing. In the cell type split, models were trained on all sequences from 80% of cell types and evaluated on the remaining 20%. Sequence-level mean PSI values were computed as the average predicted PSI across all cell types.

### Development of the Melange Model

#### Model architecture

We designed a Variational Autoencoder (VAE) based on Transformer architectures to reconstruct DNA sequences. The VAE consisted of an encoder, a latent sampling module, and a decoder. The encoder projected the one-hot encoded sequences (dimensions: batch size × 4 × 250) into a hidden representation using an initial linear projection layer. Positional embeddings were added to this projected representation to retain the positional information of nucleotides. The combined embeddings were subsequently passed through a stack of three transformer encoder layers, each consisting of multi-head self-attention with eight attention heads and feed-forward networks.

This architecture facilitated the model’s capacity to capture complex long-range dependencies within DNA sequences. The transformer encoder’s output was used to compute mean (*μ*) and log-variance (log σ2) parameters for each position in the sequence, defining a Gaussian distribution in the latent space. The latent vectors were sampled from this distribution using the reparameterization trick:

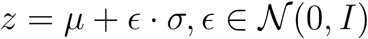

where z denotes the latent vector, *μ* and *σ* are the mean and standard deviation, respectively, and ɛ is sampled from a standard normal distribution. The sampled latent representations were decoded using a single-layer transformer encoder, aiming to reconstruct the input sequences from the latent space. The decoded outputs were projected back to the nucleotide space via a linear projection layer, yielding logits representing probabilities of nucleotide identities at each sequence position. These logits were subsequently used to reconstruct sequences through a sigmoid activation function during evaluation.

To ensure robust performance and reduce variance, we independently trained 10 Soma teacher models, each initialized with a unique random seed. Nine teachers were employed during the VAE training process to guide sequence reconstruction towards biologically meaningful PSI values. The remaining one was reserved exclusively for evaluating the biological relevance and reconstruction fidelity of sequences generated by the trained VAE.

#### Loss Function and Training Procedure

The VAE was trained using a composite loss function consisting of three components:

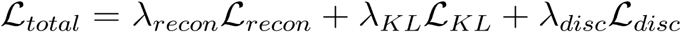

● Reconstruction Loss 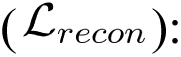 Binary Cross-Entropy (BCE) loss between the original (*x*) and reconstructed sequences 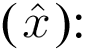

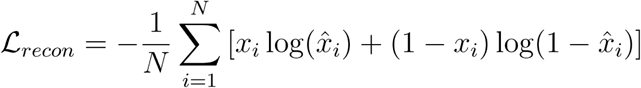

● KL Divergence Loss 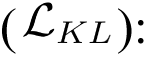 Regularization term encouraging the latent distribution to approximate a Gaussian distribution:

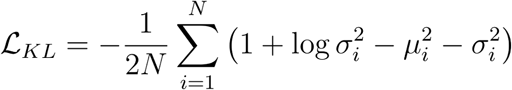

● Teacher-guided Loss 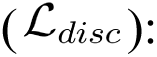 Mean Squared Error (MSE) between the teacher-predicted PSI values of reconstructed sequences 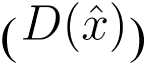 and a predefined target PSI value 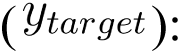

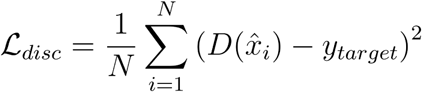

Here, 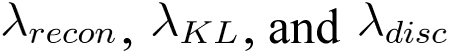 represent hyperparameters balancing the importance of each loss component.

Training utilized an Adam optimizer with a learning rate of 1e-4, batch size of 128, and was executed for 10 epochs. Dynamic weighting was employed to balance reconstruction quality and biological relevance, with hyperparameters tuned empirically.

### Design of PSI bin-switching and KELLY-specific libraries

For the PSI bin-switching library (Supplementary Table 7), we first identified sequences with stable PSI values across all cell types. Sequences were stratified into three bins: low (PSI < 0.2), intermediate (0.2–0.8), and high (> 0.8). From each bin, 500 sequences were randomly selected as starting points for Melange-designed generation. For each bin-switching task, eight designed variants were generated per parent sequence. Variants derived from parents with poor prediction accuracy were discarded, yielding a final set of 1,408 parent sequences and 5,914 designed sequences across all bins. Each library element was assigned a unique barcode and cloned using the same protocol described above.

To generate KELLY-specific sequences (Supplementary Table 8), we first defined ΔPSI as the difference between the PSI value measured in KELLY cells and the mean PSI across all other cell types. We augmented the training dataset by incorporating KELLY mutagenesis libraries, thereby expanding the sequence space and improving model robustness. We trained ten independent Soma teacher models to predict ΔPSI from input sequences. Each Soma teacher was initialized with a different random seed. For each teacher, data were randomly split into 80% training and 20% testing sets, and the models were trained for 100 epochs, during which the training loss was observed to converge. To guide the generative model (Melange), we used nine of the Soma teachers as evaluators, averaging their ΔPSI predictions to provide a stable supervisory signal. The generator was trained to produce sequences whose predicted ΔPSI matched a target value, as assessed by the ensemble of Soma teachers. This objective steered the generator toward designing sequences with strong KELLY-specific splicing repression. We trained the generator for 10 epochs for each bin-switching task. To encourage stochastic variation among sequences derived from the same parent, we applied a Kullback-Leibler (KL) divergence regularization term with a weighting factor 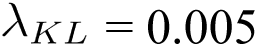 during training. The reconstruction loss, classifier guidance loss, and KL regularization were combined with weighting coefficients 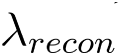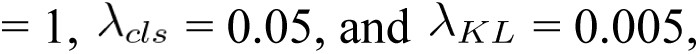 respectively. We found that setting 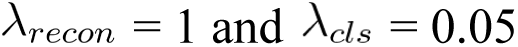 led to a reasonable number of mutation sites per sequence, while introducing a small KL regularization term 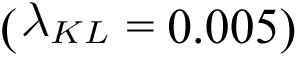 stabilized the training process.

A total of 208 parent sequences were randomly selected, and ten designed variants were generated based on each parent sequence. After filtering based on prediction accuracy, the library comprised 2,067 designed sequences. Each designed element, along with its parent sequence, was assigned a unique barcode and cloned using the same protocol described above.

### Identification of KELLY-specific sequences from designed libraries

The designed KELLY-specific library was only experimentally tested in four cell lines. To identify sequences that are KELLY-specific, we used rMATS-STAT (https://github.com/Xinglab/rMATS-STAT) to identify events that are significantly differentially spliced between KELLY and non-KELLY cells. Sequences with ΔPSI > 0.1 (group one - group two) and FDR < 0.05 were called as significant.

### Design of CRISPR guides for splicing factor mutant cell line generation

The guide sequences used to generate CRISPR-Cas9 knock-out cell lines are noted in Supplementary Table 9. All sgRNAs were validated using CRISPR amplicon sequencing of the targeted locus, and CRISPResso2 ^52^ was utilized to quantify the amount of editing (Supplementary Fig. 6a). Overexpression of splicing factor hot spot mutations was generated by cloning *U2AF1* or *SRSF2* into the plx307 backbone. Oncogenic hotspot mutations were generated using site-directed mutagenesis to introduce S34F mutants into the *U2AF1* ORF and P95H mutations into the *SRSF2* ORF.

### Generation of splicing factor mutant cells

Loss-of-function mutations in *RBM5*, *RBM10*, *FUBP1*, and *ZRSR2* were introduced into HEK293 Cas9-expressing cells transduced with sgRNAs targeting these genes and selected with puromycin at 1 μg/mL. K562 cells were transduced with sgRNAs targeting these genes selected with puromycin at 2 μg/mL. 72–96 hours after selection, K562 cells were electroporated with Alt-R CRISPR Cas9 protein (IDT). Loss of expression was validated with western blotting and amplicon sequencing of the sgRNA-targeted locus. CRISPResso2 was used to quantify indels resulting in frameshift mutations. K562 cells endogenously expressing *SF3B1*^K700E^ or wild type (*SF3B1*^K700K^) were purchased from Horizon Discovery and cultured in neomycin for selection.

HEK293 and K562 cells were transduced with lentiviral expression constructs to overexpress wild-type or mutant forms of *U2AF1* and *SRSF2*. Selection with puromycin (1 μg/mL or 2 μg/mL) was performed before validation of expression by western blotting for a C-terminal V5 tag.

### Western Blot Analysis

Cell pellets were lysed in RIPA buffer supplemented with protease inhibitor cocktail (Thermo Scientific) and incubated on ice for 15 minutes. Lysis was facilitated by vortexing every five minutes before the pellet was separated from the supernatant by centrifugation at 15000 rpm for 15 minutes at 4°C. Total protein amounts were quantified using the Bicinchoninic Acid (BCA) assay, and samples were denatured in NuPage sample buffer containing BME and boiled at 95°C for 10 minutes. Approximately 20–30 μg of protein were loaded into each well of a 4–12% gradient Bis-Tris gel and resolved by SDS-PAGE before being transferred to a nitrocellulose membrane via wet transfer for 2 hours at 4°C. Membranes were blocked with Intercept (PBS) blocking buffer and blotted overnight with primary antibodies directed to RBM5 (Cell Signaling, #86425), RBM10 (Cell Signaling 18012S), FUBP1 (Sigma ABE1330), V5 (Cell Signaling D3H8Q), Vinculin (Millipore sigma V9131), or beta-actin (Cell Signaling 8H10D10).

Membranes were washed and probed with fluorescently tagged secondary antibodies (Licor), and visualized on the Licor Odyssey M.

### Design of splicing factor mutation-specific designed library

To generate sequences with splicing activity biased toward either wild-type (WT) or mutant (MUT) cellular backgrounds, we defined ΔPSI as the difference in splicing ratios between the two conditions, calculated as PSI(WT) - PSI(MUT). Sequences with ΔPSI values close to zero were selected as “neutral” backbones for targeted sequence editing. We applied the same training and generation framework as described for the KELLY analysis. We trained 10 Soma models to estimate ΔPSI from sequence inputs, and their averaged predictions were used to provide guidance during Melange training. Melange was trained to reconstruct input sequences while being optimized to generate sequences whose predicted ΔPSI matched a target value. The loss function combined reconstruction loss, classifier guidance loss, and KL regularization with weighting coefficients 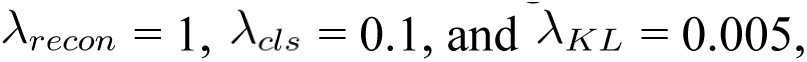 respectively. We found that setting 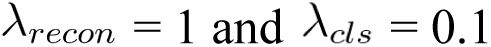 produced a reasonable number of mutations per sequence, while the small KL regularization 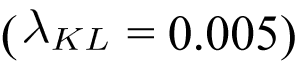 stabilized training. Melange was trained for 20 epochs, effectively directing mutations toward sequences predicted to exhibit strong differential splicing activity between WT and MUT cells. We randomly selected 581 parent sequences that were not differentially spliced between WT and MUT cells, and designed eight variants per parent sequence (Supplementary Table 10). After filtering, the final libraries comprised 1,734 sequences for *RBM5* knockout and 440 sequences for *RBM10* knockout. Each designed element, together with its parent sequence, was assigned a unique barcode and cloned using the same protocol described above.

### Identification of splicing-factor mutant-specific sequences

The designed splicing-factor mutant-specific library was experimentally tested in pairs of cell lines (WT and mutant), so again we used rMATs to calculate p-values and FDRs (false discovery rates) between the two groups. Sequences with ΔPSI > 0.1 (group one - group two) and FDR < 0.05 were called as significant, leading to a total of 5 significant sequences in the *RBM5* KO cells and 16 significant sequences in the *RBM10* KO cells.

## Acknowledgements

We thank members of the Chen lab and Golub lab for helpful discussions. We would also like to thank P. Sharp, A. Krishnan, V. Viswanadham, S. Rong, and J. Roth for helpful discussions. We also thank A. Patentreger and A. Masse at the Broad Institute Flow Cytometry Core Facility for their experimental support. The authors used ChatGPT to improve readability and language clarity. The authors take full responsibility for the content.

## Authors contributions

Conceptualization, X.D.C. and F.C.; Methodology, X.D.C., M.J., M.V., and K.C.; Investigation, X.D.C., M.J., M.V., A.N.T., J.W.L., Y.Z., D.W., J.S., G.S., S.M., M.H., Software, X.D.C., M.J.

Q.G., and K.C.; Data curation, X.D.C. and M.J.; Formal Analysis, X.D.C., M.J., and M.V.; Visualization, X.D.C., M.J., M.V., and K.C.; Resources, J.R.; Writing - Original Draft, X.D.C., M.J., M.V., and K.C.; Writing - Review and Editing, J.W.L. and F.C; Funding Acquisition, T.R.G. and F.C., Supervision, T.R.G. and F.C..

## Data and materials availability

Sequencing data will be deposited in the Gene Expression Omnibus (GEO, accession number TBD). Plasmid backbones generated for this study will be deposited on AddGene. Code used to process data and generate figures is located at https://github.com/chen-dawn/SPICE. Packages for machine learning models are located at https://github.com/caokai1073/SPICE-ML.

---

**Supplementary Figure 1:**
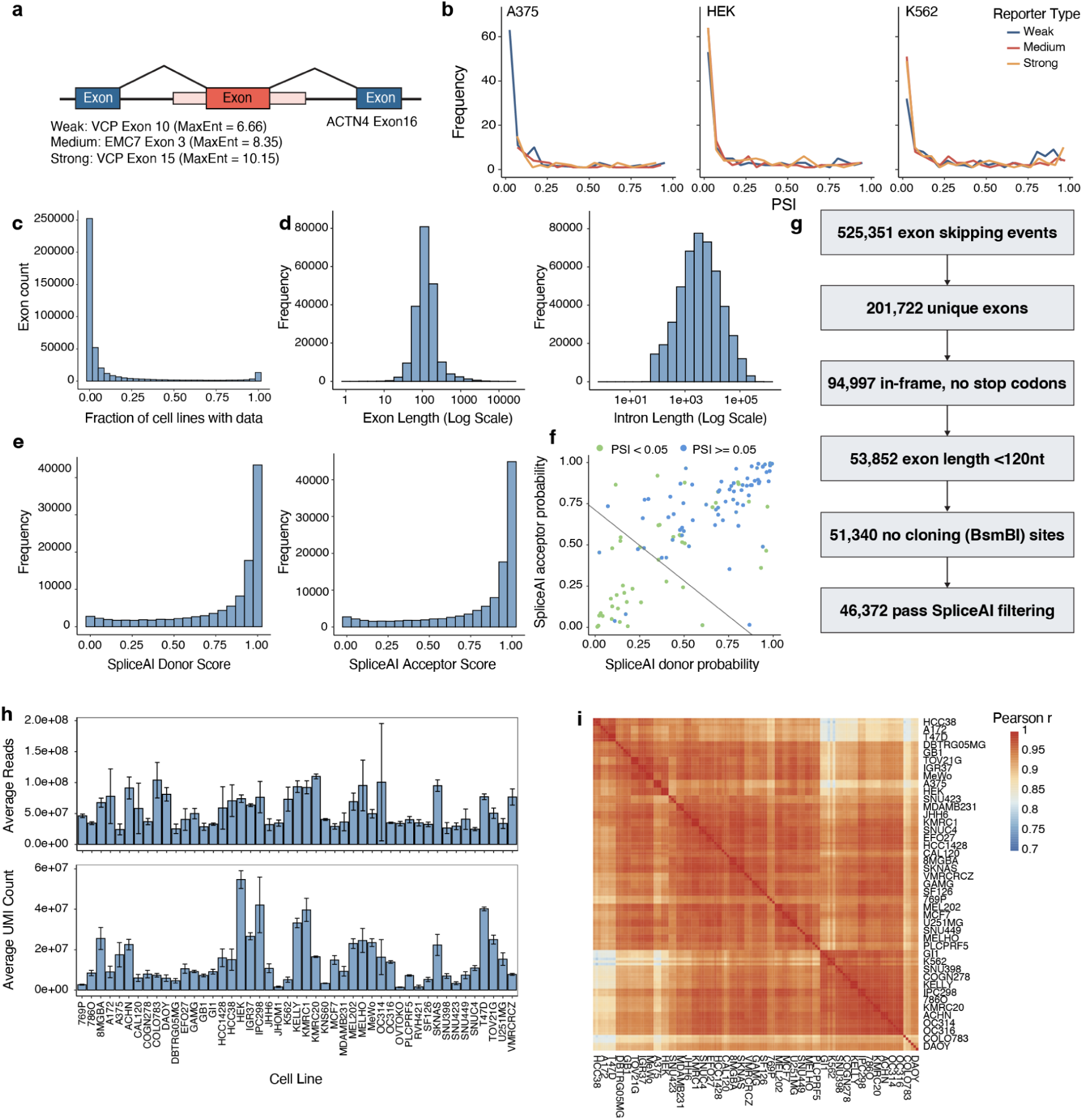
Reporter design, library construction, and data quality control. **a**, Exon skipping reporter constructs with weak (*VCP* Exon 10, MaxEntScan = 6.66), medium (*EMC7* Exon 3, MaxEntScan = 8.35), or strong (*VCP* Exon 15, MaxEntScan = 10.15) 5′ splice sites. All constructs contain *ACTN4* Exon 16 as the constant downstream exon. **b**, PSI distributions across weak, medium, and strong reporter backbones in A375, HEK293, and K562. **c**, Fraction of CCLE cell lines with quantifiable exon skipping events from RNA-seq. **d**, Exon length (left) and intron length (right) distributions for candidate library elements. **e**, Distribution of SpliceAI-predicted donor (left) and acceptor (right) scores for candidate exons. **f**, Scatterplot of SpliceAI donor and acceptor probabilities, colored by PSI measured in pilot experiments. **g**, Filtering workflow for construction of the final exon library. **h**, Average reads (top) and unique molecular identifier (UMI) counts (bottom) per cell line. Error bars show standard deviation across replicates. **i**, Pairwise Pearson’s correlation coefficients of exon PSI values between replicates across all profiled cell lines.

**Supplementary Figure 2:**
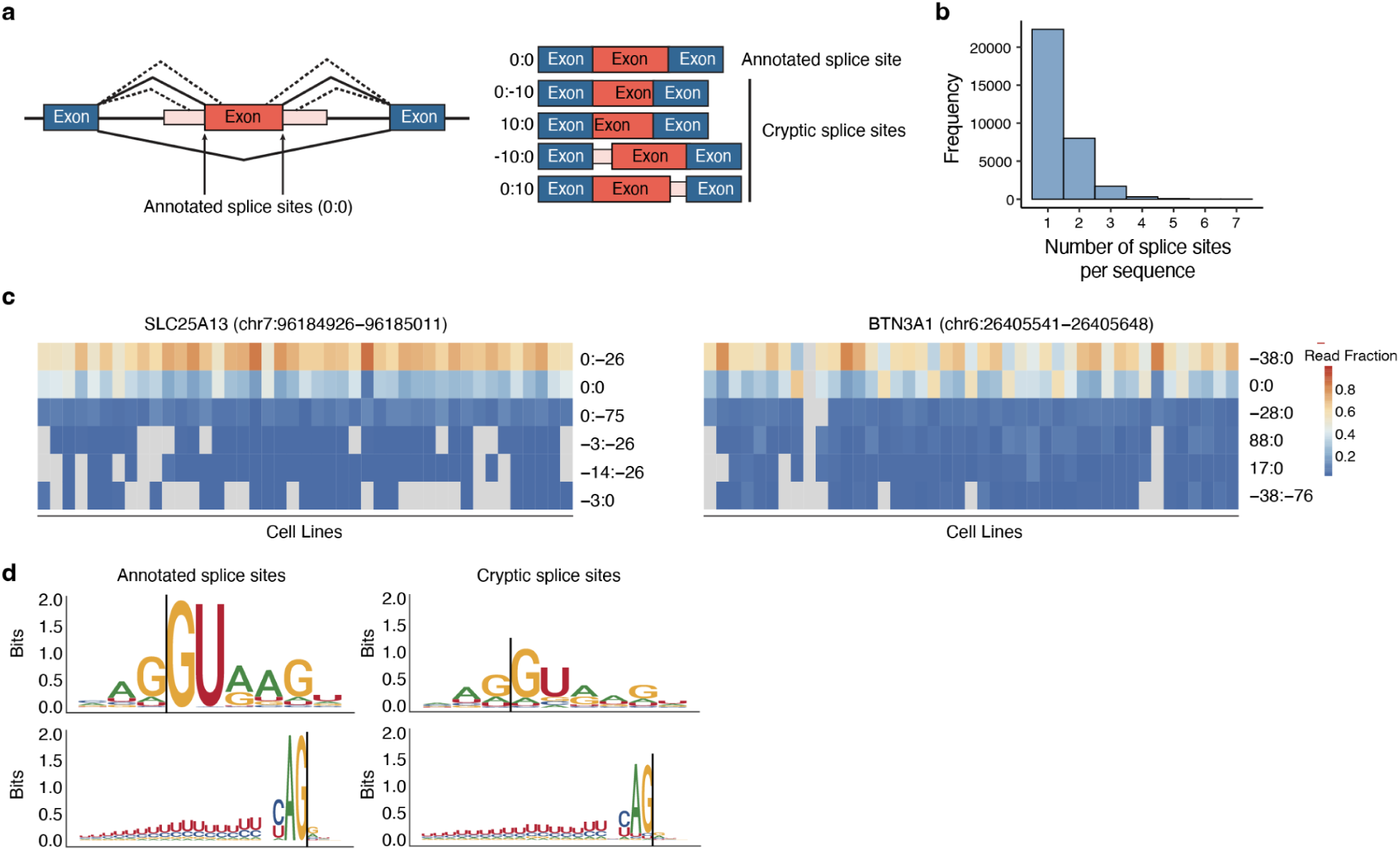
Cryptic and annotated splice site usage across library elements. **a**, Schematic of annotated versus cryptic splice site usage within the exon skipping reporter system. Cryptic splice sites use non-annotated 3′ or 5′ sites, and can result in additional or fewer nucleotides at each end of the library exon. **b**, Histogram of the number of splice sites detected per library element. **c**, Representative examples of library elements with multiple splice sites used consistently across cell lines, showing annotated and cryptic splice site junction usage. **d**, Sequence logos of annotated donor (5′, top left) and acceptor (3′, top right) splice sites, and cryptic donor (5′, bottom left) and acceptor (3′, bottom right) splice sites.

**Supplementary Figure 3:**
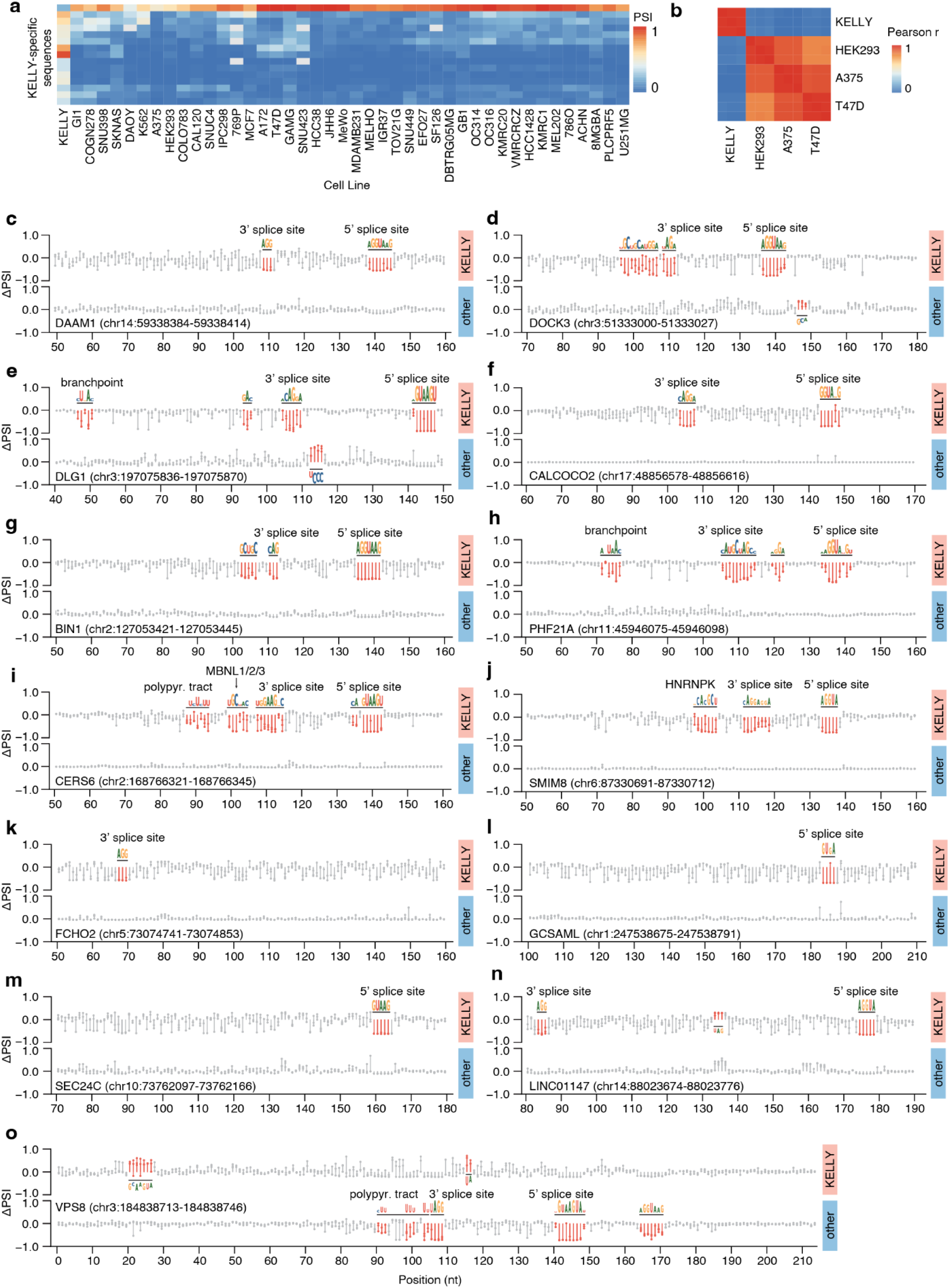
Saturation mutagenesis reveals sequence determinants driving cell type-specific splicing. **a**, Heatmap of fifteen KELLY-specific elements across all cell lines from the unbiased MPRA library, selected to use for saturation mutagenesis. Heatmap color scale indicates PSI, where red represents PSI = 1, and blue represents PSI = 0. **b**, Heatmap of Pearson’s correlations of PSI between cell lines profiled in the saturation mutagenesis library. **c-o**, Effect size ΔPSI for all saturation mutagenesis variants, separated by parent sequence and between KELLY and non-KELLY cells. Sequence motifs are splicing sensitive regions called by our *z*-score filtering approach (Methods). **c**, *DAAM1* (chr14:59338384–59338414) **d**, *DOCK3* (chr3:51333000–51333027) **e**, *DLG1* (chr3:197075836–197075870) **f**, *CALCOCO2* (chr17:48856578–48856616) **g**, *BIN1* (chr2:127053421–127053445) **h**, *PHF21A* (chr11:45946075–45946098) **i**, *CERS6* (chr2:168766321–168766345) **j**, *SMIM8* (chr6:87330691–87330712) **k**, *FCHO2* (chr5:73074741–73074853) **l**, *GCSAML* (chr1:247538675–247538791) **m**, *SEC24C* (chr10:73762097–73762166) **n**, *LINC01147* (chr14:88023674–88023776) **o**, *VPS8* (chr3:184838713–184838746) Polypyr. tract: polypyrimidine tract.

**Supplementary Figure 4:**
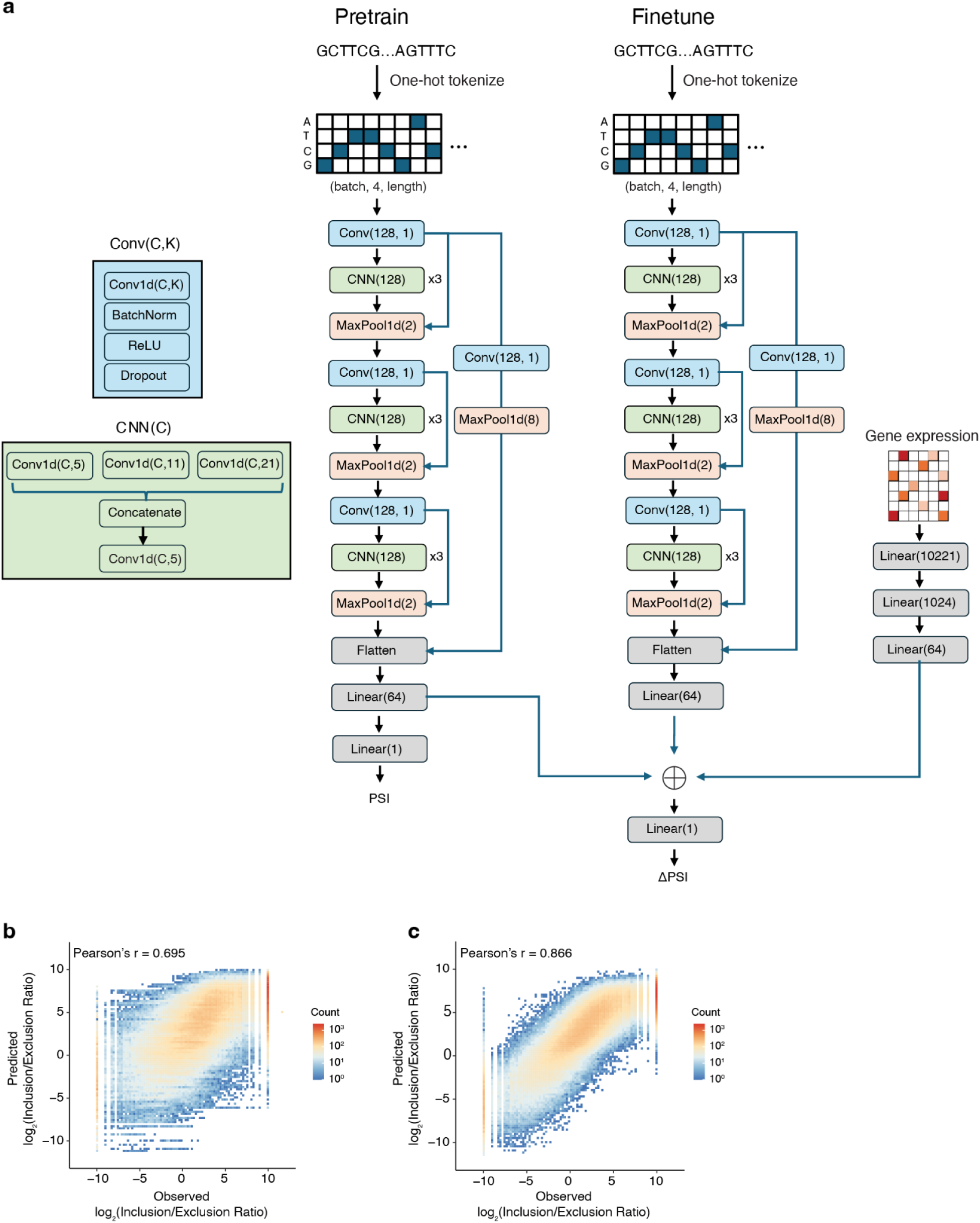
Soma model structure and prediction accuracy. **a**, Schematic of the Soma model architecture. **b,** Scatter of Pearson’s correlations between predicted and observed splicing outcomes for a held-out set for unseen sequences using only sequence information. **c,** Scatter of Pearson’s correlations between predicted and observed splicing outcomes for a held-out set for unseen cell types using only sequence information.

**Supplementary Figure 5:**
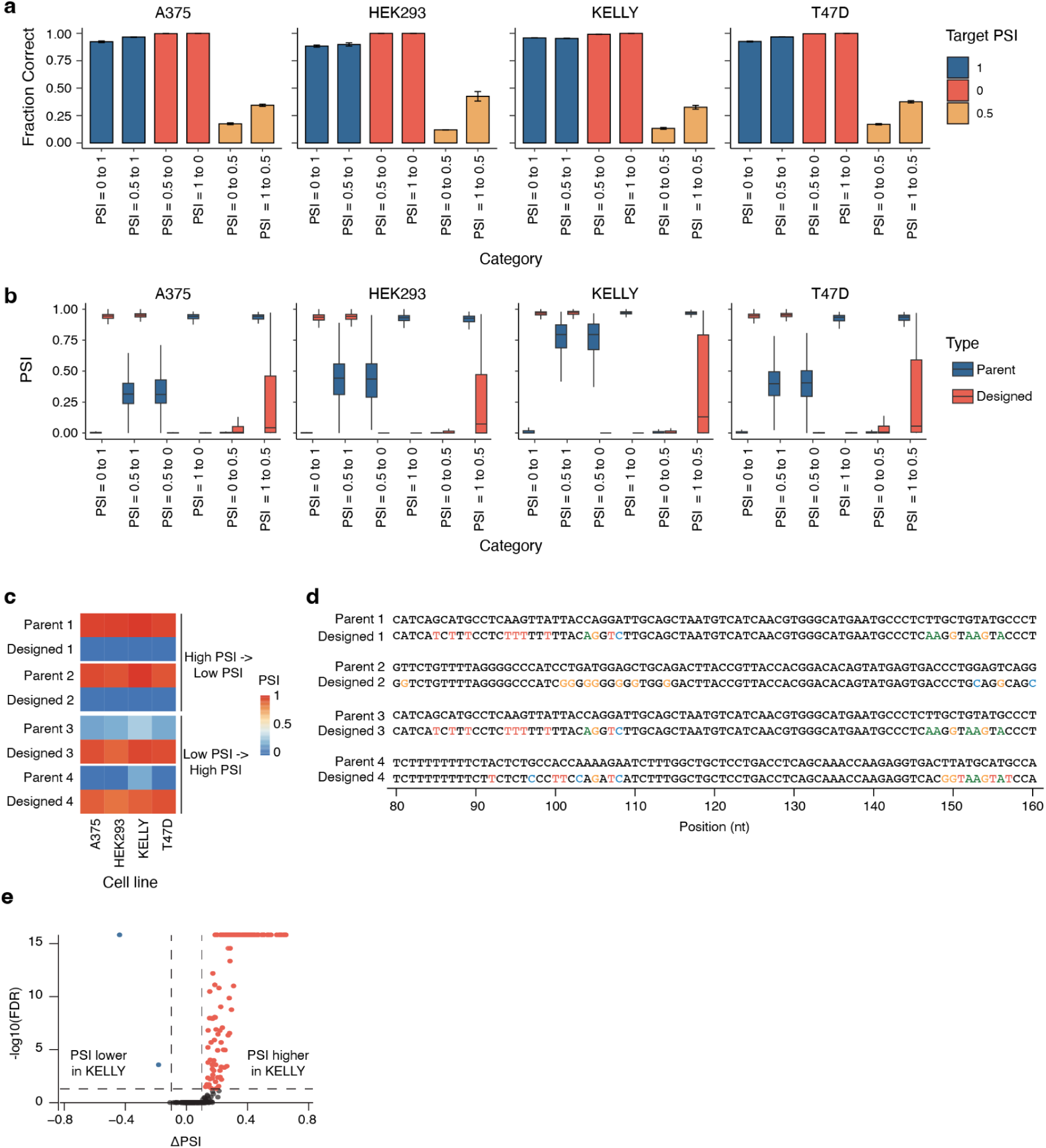
Melange accurately generates sequences with user-specified splicing profiles. a, Fraction of sequences achieving the intended outcome across design categories in the profiled cell lines. Error bars indicate standard deviation across experimental replicates (n = 3). **b**, Distribution of PSI values for designed sequences across target categories in the profiled cell lines. **c**, PSI values of example sequences from the bin-switching task, for both the parent and designed sequences in the categories {high PSI (>0.8) to low PSI (<0.2)} and {low PSI (<0.2) to high PSI (>0.8)}. Values are averaged across three replicates in each cell line. **d**, Example sequences from the bin-switching task. Highlighted nucleotides indicate differences between the parent sequence and the designed sequence. **e**, Differential splicing of KELLY-specific designed sequences identified by rMATS, comparing PSI between KELLY and non-KELLY cells. Significant events were defined as ΔPSI > 0.1 and FDR < 0.05. Red points indicate higher PSI in KELLY cells, and blue points indicate lower PSI in KELLY cells.

**Supplementary Figure 6:**
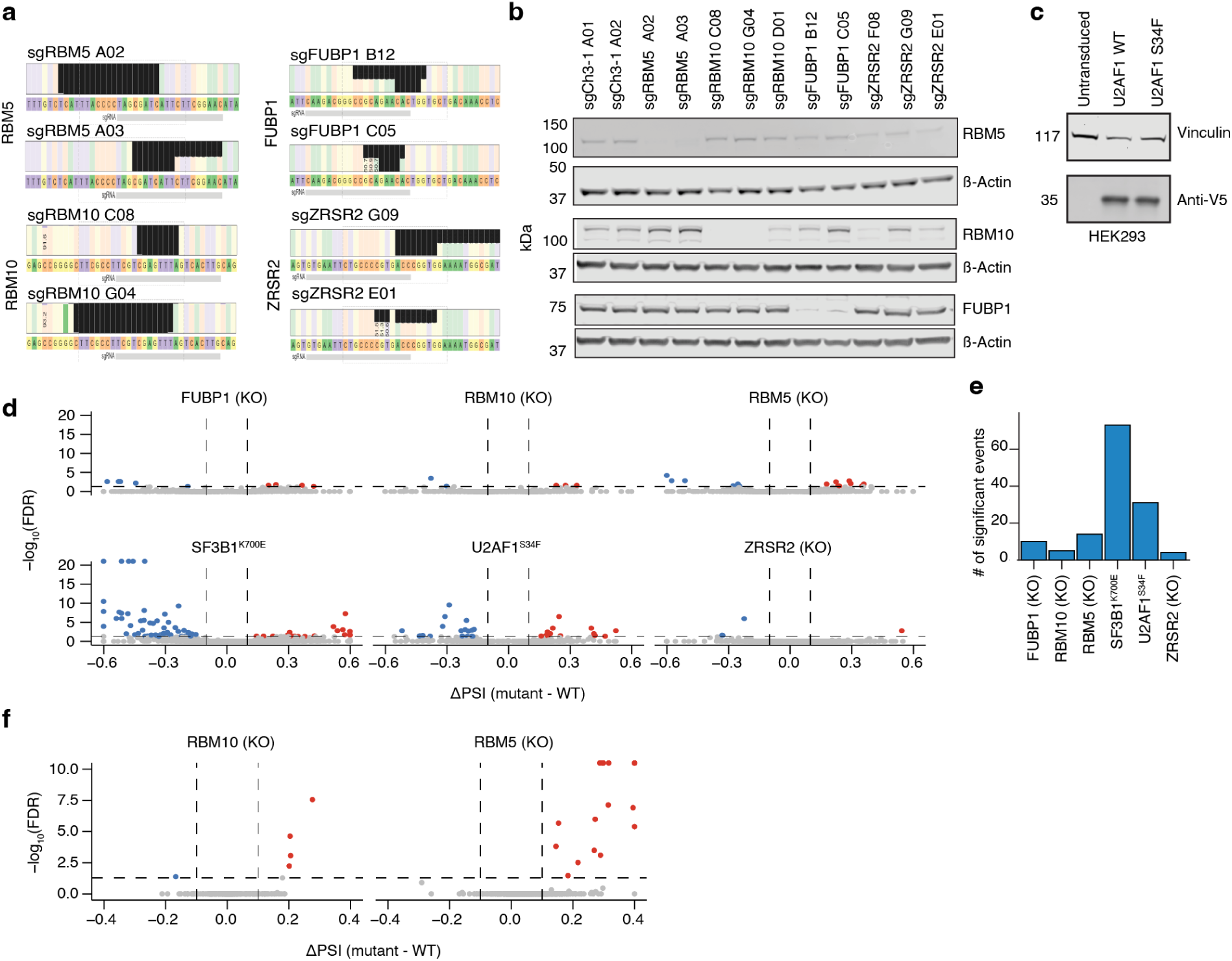
Cells harboring cancer mutations display misregulated splicing profiles. **a**, CRISPR knock-out sites for K562 single cell clones, for *RBM5* KO, *RBM10* KO, *FUBP1* KO, and *ZRSR2* KO cells. **b**, Western blotting for RMB5, RBM10, and FUBP1 in each of the single cell K562 clones. Ch3-1 guides are non-targeting sgRNAs. ZRSR2 expression is not shown as no commercial antibodies had sufficient binding affinity. **c**, Western blotting for V5-tagged U2AF1^WT^ and U2AF1^S34F^ expression in HEK293. Cells were infected with lentivirus containing a transgene with the corresponding *U2AF1* cDNA sequence. **d**, Volcano plots of significant library elements identified per mutant background, for alternative 3′ splice site selection. Significant events have ΔPSI > 0.1 and FDR < 0.05. Significant points in red have higher PSI values in the KO cells than WT cells, and points in blue have lower PSI values in the KO cells than WT cells. **e**, Number of significant library elements identified per mutant background, filtered by ΔPSI > 0.1 and FDR < 0.05, for alternative 3′ splice site selection. **f**, Volcano plots of significant sequences between mutant and WT paired cell lines in alternative 3′ splice site selection. Significant events have ΔPSI > 0.1 and FDR < 0.05. Significant points in red have higher PSI values in the KO cells than WT cells, and points in blue have lower PSI values in the KO cells than WT cells. KO: knock-out. WT: wild-type.

