## Supplementary Note 1 for "Generative Design of Cell Type-Specific RNA Splicing Elements for Programmable Gene Regulation"

### Supplementary Note 1: Evaluating cell line transfection efficiency across 62 cell lines

#### Cell line pooling and transfection

Before performing the individual MPRA experiments, we set out to determine the best transfection condition for each cell line. Because we obtained our cells from the PRISM assay[1], each cell line contains individual DNA barcodes, enabling transfection condition optimization in a pooled approach[2].

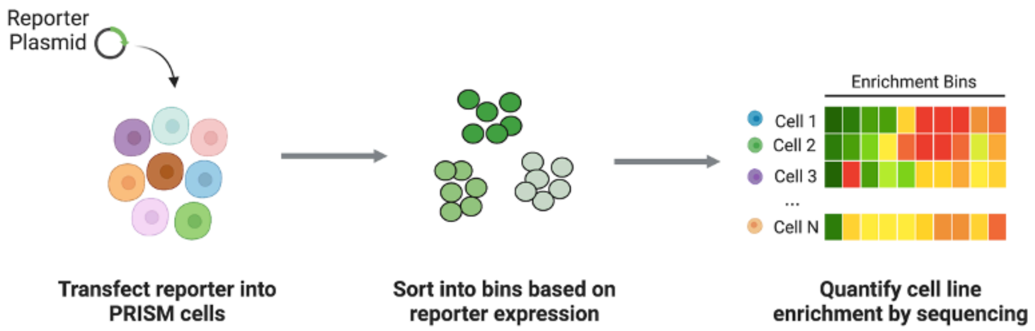


**Figure 1**: Schematic of pooled optimization protocol.

We employed PRISM barcoded cell lines reflecting both solid and hematopoietic lineages to determine transfection efficiency using both electroporation and chemical transfection approaches (**Figure 1**). Cells were individually grown, then pooled into 2 pools – one pool for adherent cells (non-hematopoietic lineage) and one pool for suspension cells (hematopoietic lineage) – on the day of the experiment. Each cell pool was transfected with a 7.5 kb plasmid containing a mCherry fluorescent reporter. We tested 6 different electroporation methods, which involved different combinations of pulse codes and nucleofection buffer on the 4D-LonzaX Nucleofector. These conditions included: CM137 (P2, P5, SE, SF buffer), DS137 (SF buffer), EN137 (SF buffer), FF120 (SF buffer), CM137 (SG buffer). Cells were also chemically transfected with TransIT-2020 (liposomal transfection reagent) or TransIT-X2 (non-liposomal polymeric transfection reagent) from Mirus Bio. 2 million pooled cells were transfected for each condition in duplicate.

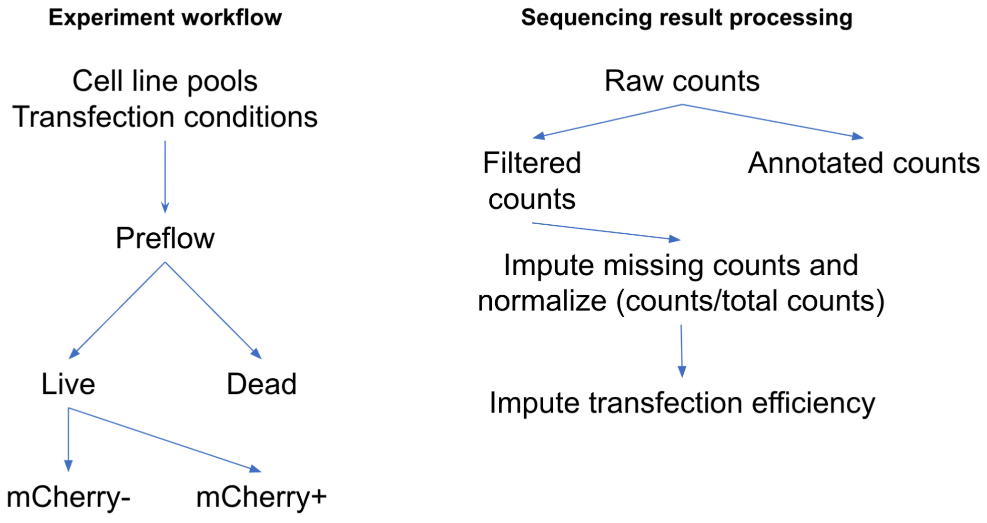


**Figure 2**: Schematic of flow sorting and sequencing data analysis. In the experiment workflow, cell samples were collected for each of the populations: preflow, live, mCherry-, mCherry+.

**FACS Sorting**

24-48 hours following transfection, cell line pools were harvested using trypsinization for adherent cells. Samples were washed and resuspended in flow buffer (PBS with 2% FBS and 0.5 mM EDTA) and stained with DAPI viability dye. Pooled cells that did not undergo transfection were used to establish a negative gate for flow sorting. Sorting was performed on SONY SH800 or MA900 sorters. Dead cells (DAPI-positive cells) were sorted into a separate bin from live (DAPI-negative cells). Viable cells (DAPI negative) were sorted into mCherry-positive and mCherry-negative bins (**Figure 2**). The PRISM barcodes in these sorted populations were sequenced to determine the transfection efficiency. Barcode abundance was compared to barcode abundance from the pre-sort population.

**Sequencing of PRISM barcodes**

Samples were directly lysed in DNA Lysis Buffer (20mM Tris-HCl, 50mM KCl, 0.45% NP-40, 0.45% Tween-20, 10% Proteinase K) and denatured at 95 ºC and amplified with a Q5 polymerase master mix. PRISM sequencing primers allowed samples to be dual-indexed for multiplexed Illumina sequencing by directly adding Illumina flow-cell binding sequences to the amplicon:

forward primer:
5’AATGATACGGCGACCACCGAGATCTACANNNNNNNNAAGGTGCTTCTCGATCTGCAT
reverse primer:
5’CAAGCAGAAGACGGCATACGAGATNNNNNNNNGTGACTGGAGTTCAGACGTGTGCT.

where N represents the indexing nucleotides. Resulting products were evaluated for single-band amplification using gel electrophoresis before being pooled and purified for sequencing using the Zymo Select-a-Size DNA Clean & Concentrator kit. After pooling, the PCR product was quantified using the Qubit 3 Fluorometer and the size was estimated using an Agilent Tapestation. Samples were sequenced using Illumina NextSeq technology. Samples were loaded onto the NextSeq flow cell at a final concentration of 10 pM with a 20% PhiX spike-in due to low diversity. Sequencing was run for 50 cycles, single-read.

$$cellline fraction_{mCherry} = \frac{\left[\frac{barcode_{PRISM}\, counts}{totalcounts}\right]_{mCherry+} \cdot pool_{mCherry+}}{\left\{\left[\frac{barcode_{PRISM}\, counts}{totalcounts}\right]_{mCherry+} \cdot pool_{mCherry+}\right\} + \left\{\left[\frac{barcode_{PRISM} counts}{totalcounts}\right]_{mCherry-} \cdot pool_{mCherry-}\right\}}$$

$barcode_{PRISM}$ counts = # cell line reads in mCherry+/- population sequenced
total counts = # PRISM reads in mCherry+/- population sequenced
$pool_{mCherry+}$= fraction of total pool sorted into mCherry+ bin
$pool_{mCherry-}$= fraction of total pool sorted into mCherry- bin

**Figure 3**: Formula for normalizing counts to determine cell line transfection efficiency.

**Quantification of transfection efficiency**

To infer the transfection efficiency of individual cell lines for each condition based on the pooled sequencing read counts, we used the formula described in **Figure 3**. This analysis gave us the relative transfection efficiency of all cell lines profiled (**Figure 4a-b**). To validate the pooled results, we benchmarked transfection efficiency in eight individual cell lines using the 4D-Lonza X Nucleofector with the SF buffer and a range of program codes. Transfection efficiency in these single-cell-line experiments was quantified via individual flow cytometry analysis (**Figure 4c**). Although the absolute efficiency values derived from pooled sequencing and individual FACS measurements did not match perfectly, their trends were highly correlated, indicating that pooled results provide a reliable proxy for relative transfection efficiency across conditions.

Overall, we found that chemical transfection reagents performed better than electroporation in non-hematopoietic (adherent) cell lines, whereas hematopoietic (suspension) cell lines were transfected poorly by chemical-based methods and can only be electroporated. Cell lines that were transfectable generally exhibited good efficiency across multiple conditions, while certain lineages remained difficult to transfect regardless of approach. Based on these results, we selected chemical transfection reagents for non-hematopoietic cell lines in subsequent MPRA experiments.

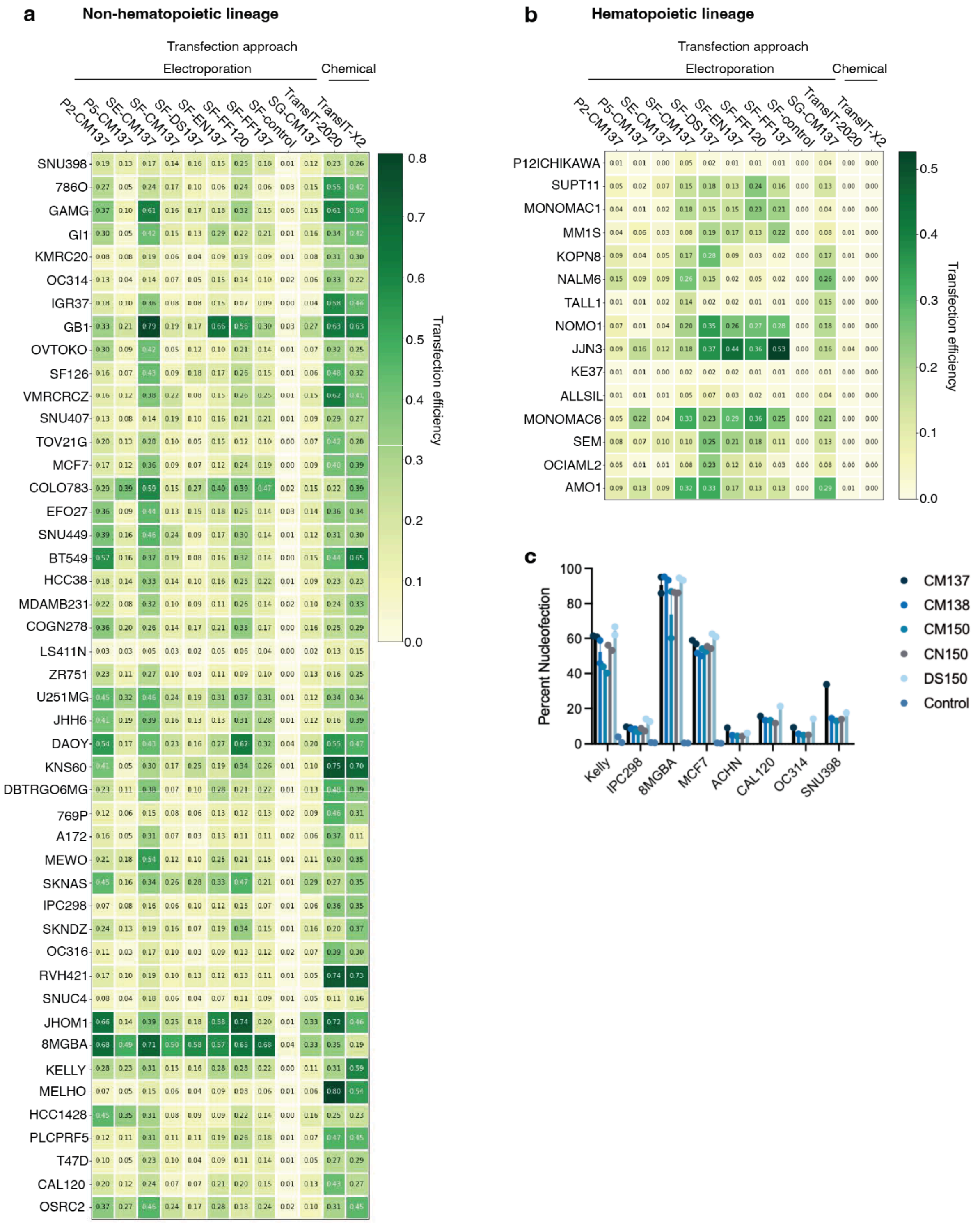


**Figure 4** (a) Relative transfection efficiency in adherent cell lines of non-hematopoietic origin. (b) Relative transfection efficiency in hematopoietic cell lines. (c) Transfection efficiency for single cell line benchmarks.
