## Supplementary Note 2 for "Generative Design of Cell Type-Specific RNA Splicing Elements for Programmable Gene Regulation"

### Supplementary Note 2: Correcting for Batch Effects Due to Barcode Swapping

We observe a phenomenon we term *PSI depression*, where sequences that should be constitutively included exhibit unexpectedly lower inclusion levels (for example, a sequence expected to have PSI = 1 may show a measured PSI = 0.95). This effect arises from barcode swapping during PCR amplification, leading to the generation of chimeric reads in which the barcode from one molecule is incorrectly transferred to another.

Recall that the percentage splice inclusion (PSI) is calculated as:

$$\text{PSI} = \frac{I}{I+E}$$

where $I$ and $E$ denote the number of reads corresponding to the included and exon-skipped isoforms, respectively.

In our sequencing-based readout, *Read 1* spans exon–exon junctions and defines the splicing state of the sequence, while *Read 2* identifies the library element barcode. When the middle exon is included, *Read 1* and *Read 2* can be cross-referenced to verify concordance and detect barcode swapping events, allowing us to estimate the chimeric rate, $C$. However, when the exon is skipped, *Read 1* provides no information about the original molecule's identity. In this case, read assignment relies solely on the barcode, which makes skipped isoforms particularly vulnerable to misassignment caused by barcode swapping.

Ideally, barcode swapping should be minimized experimentally. We have implemented several optimizations to reduce the swapping rate, including decreasing the number of PCR cycles and extending the elongation time per cycle, as described by Omelina et al., 2019. Although these adjustments help reduce the formation of chimeric DNA molecules, residual artifacts remain and require computational correction post-sequencing.

**Computational Correction Model**

To perform the correction, we incorporate both included and skipped reads into the correction model.

Let $I$ denote the number of included reads, $E$ the number of exon-skipped reads, and $C$ the chimeric rate. The chimeric rate, $C$, is empirically estimated by measuring the concordance between *Read 1* and *Read 2* in the exon-included reads population for each sample. The total number of reads is defined as $T = I + E$. The observed number of exon-skipped reads is then given by:

$$\hat{E} = E + C(I+E)$$

Solving for the true number of skipped reads $E$:

$$E = \frac{\hat{E} - CI}{1+C}$$

In the case of multiple exon-skipped categories $(E_1, E_2, \ldots, E_n)$, we generalize the model as follows. The observed total of skipped reads is:

$$\hat{E} = (E_1 + E_2 + \ldots + E_n) + C(I + E_1 + E_2 + \ldots + E_n)$$

Solving for the true total skipped reads:

$$E_1 + E_2 + \ldots + E_n = \frac{(\hat{E}_1 + \hat{E}_2 + \ldots + \hat{E}_n) - CI}{1+C}$$

Finally, for each element $E_i$, the corrected value can be computed proportionally as:

$$E_i = \frac{\hat{E}_i}{\sum_{j=1}^{n} \hat{E}_j} \times \frac{\sum_{j=1}^{n} \hat{E}_j - CI}{1+C}$$
